# Inferring ecDNA Evolution from Allele Frequency Spectra

**DOI:** 10.64898/2026.10.09.758054

**Authors:** Bhargavi Dameracharla, Gino Prasad, Jens Luebeck, Samir Amin, Jonathan Kirkland, Owen S. Chapman, Lukas Chavez, Roel Verhaak, Weini Huang, Paul S. Mischel, Ben Werner, Vineet Bafna

## Abstract

Extrachromosomal DNA (ecDNA) can arise early in the development of aggressive tumors or later upon progression, but the impact of chronology on patient outcome is not known. Currently, it is not possible to determine when an ecDNA arose in the course of tumor development and progression, based on genome sequencing data. Here, we develop an analytic method, ecEvol, to infer ecDNA emergence time, selection strength, mutation rate, and clonal dominance from allele-frequency spectra in bulk whole-genome sequencing. EcEvol outperforms other methods and predicts parameters with high accuracy across a wide range of parameter values. Applying ecEvol to primary, treatment-naive tumors, we find that ecDNA typically emerges early and undergoes strong positive selection, albeit with substantial variation across cancer types. EcDNAs carrying known oncogenes emerge earlier and reach greater clonal dominance than ecDNAs lacking known oncogenes. In advanced and predominantly treated tumors, ecDNA emerges substantially later than in matched primary tumors, consistent with the generation and selection of new ecDNA during tumor progression and therapy. Longitudinal samples further support continued expansion of ecDNA-bearing lineages. Finally, earlier ecDNA emergence is associated with poorer overall survival. These results provide insight into the chronology of ecDNA formation from a single tumor biopsy taken at any point during the course of the disease, revealing that allele-frequency spectra encode the evolutionary history of ecDNA and provide a quantitative framework for reconstructing its emergence and expansion.

## Introduction

Cancer is an evolutionary process driven by mutation and selection ^1^ and reshaped by microenvironmental selection pressures ^2^. As environmental niches change over space and time, selective pressures alter the fitness landscape, enabling specific mutations to confer growth or survival advantages that drive the expansion of distinct subclones at multiple locations and time points during the lifetime of a single tumour ^3,4^.

The fitness landscape is dramatically altered when oncogenic alterations are carried on extrachromosomal DNA (ecDNA). EcDNAs are large, circular, acentric molecules comprised of chromosomal DNA but reside and replicate outside of canonical chromosomes ^5^. Lacking centromeres, ecDNAs segregate randomly among daughter cells at mitosis. The stochastic and non-Mendelian inheritance promotes rapid oncogene amplification and extensive intratumoral copy-number heterogeneity^6^. EcDNA-mediated oncogene amplification has been detected across a broad spectrum of cancer types, is enriched in advanced and treatment-resistant tumours, and is associated with significantly shorter patient survival ^5–7^.

Theoretical models of ecDNA population dynamics, together with empirical observations, have established that random segregation during mitosis and selective growth advantage jointly shape the distribution of ecDNA copy number across tumour cells^6^. However, these models do not predict when ecDNA first emerged during tumour evolution. Empirical evidence suggests that ecDNA may arise in premalignant tissue in some contexts ^8,9^, while in others it appears much later, potentially in response to chemotherapeutic selection pressures^7,10–12^. Similarly, ecDNA copy number varies substantially across tumours, ranging from only a few copies per cell in some cancers^5^ to hundreds of copies per cell in others. These observations likely reflect differences in both the timing of ecDNA emergence and the magnitude of the selective fitness advantage conferred by ecDNA, but the relationship between these parameters has not been quantitatively characterized.

In contrast, prior work on subclonal architectures arising from chromosomal mutations has demonstrated that both selective fitness advantage and the timing of subclone emergence can be jointly inferred from variant allele frequency distributions^13^. Many methods infer tumour evolutionary parameters from the variant allele frequency distributions obtained by bulk whole-genome sequencing ^14,15^. More recently, deep learning approaches extend these ideas by training neural networks on forward simulations to recover selection coefficients and subclone emergence time from chromosomal variant allele frequency distributions in single biopsies ^16,17^. These methods were designed for chromosomally inherited subclones and do not account for the distinct allele frequency structure generated by ecDNA. Stochastic ecDNA segregation and dynamic copy-number amplification reshape the mutational spectrum of the ecDNA containing region relative to spectrum of the chromosomal mutations: intermediate-frequency variants are depleted, while a high-frequency cluster of ecDNA-amplified mutations emerges whose structure encodes the evolutionary history of the ecDNA-bearing lineage. To our knowledge, existing frameworks do not exploit this dual-spectrum structure for ecDNA-specific evolutionary inference.

Here, we develop ecEvol, a simulation and deep learning framework to infer ecDNA evolutionary parameters directly from bulk whole-genome sequencing data. Building upon theoretical models of stochastic ecDNA population dynamics, we implemented a forward simulator that explicitly models random ecDNA segregation and amplification dynamics. We then trained a cross-attention deep learning architecture to jointly infer selection strength, mutation rate, and the timing of ecDNA emergence, together with a companion regression model that estimates the fraction of tumour cells descended from the ecDNA-founding ancestor. We then apply these methods to wgs data from cancer genome projects to characterize how these parameters vary across tumour samples and tested their association with clinical and molecular features of those samples.

## Results

### The allele frequency spectrum informs the timing and strength of subclonal expansions

To recapitulate how subclones alter the allele frequency spectrum (AFS), we first examined the AFS in simulations where no subclone is present. We modeled a population of tumor cells arising at time *T*_0_ from a single founder clone carrying *C* clonal (driver and hitchhiker) mutations (Figure 1a). Tumor growth proceeded exponentially from this founder clone under a stochastic branching process, and new somatic point mutations were introduced at division events (full parameterization in Methods).

**Figure 1:**
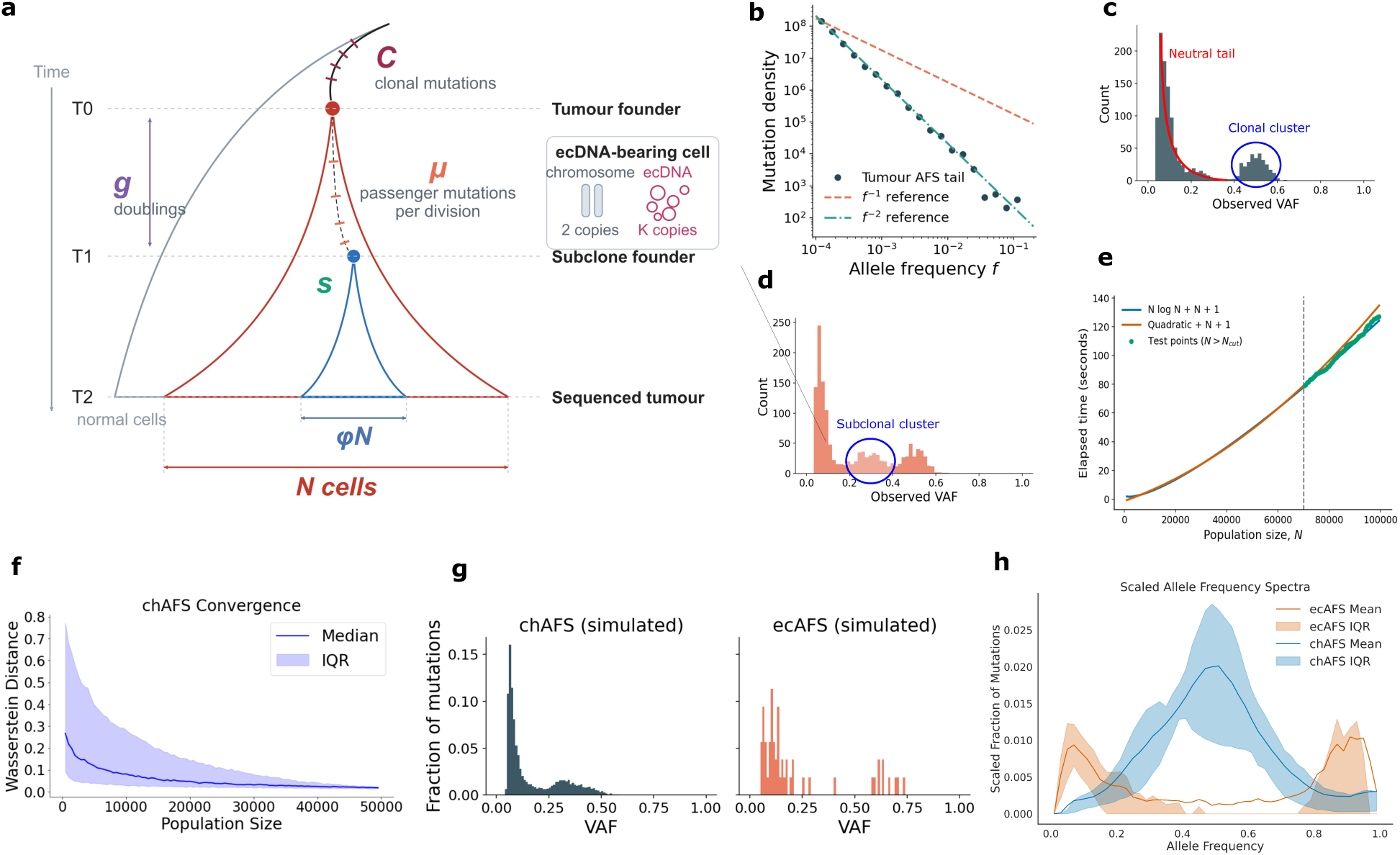
Allele frequency spectra of ecDNA(+) tumours. **(a)** Schematic of the stochastic branching process model. The tumour founder at *T*0 carries *C* clonal mutations; cells acquire *µ* passenger mutations per division; after *g* doublings a fitter subclone with selection coefficient *s* arises at *T*1 and by sampling (*T*2) accounts for a fraction *ϕ* of the *N* tumour cells. Inset: an ecDNA-bearing cell carries *K* ecDNA copies alongside two chromosomal copies of the region. **(b)** The allele frequency spectrum of a tumor grown under neutral exponential expansion exhibits the characteristic *f^−^*^2^ decay at low frequencies. Each point represents a unique observed mutation density in the population at the end of simulation. Dashed lines show *f^−^*^1^ and *f^−^*^2^ reference slopes. **(c)** Simulated bulk sequencing of a tumor containing clonal mutations produces a prominent ‘clonal peak’ centered at 0.5 in the diploid genome, with remaining passenger mutations contributing to a ‘neutral tail’ at lower frequencies. **(d)** Introduction of a fitter subclone (selection coefficient *s >* 0) generates an additional cluster of subclone–private mutations at intermediate frequency. **(e)** Comparison of *N* log *N* and quadratic fits; the *N* log *N* model generalizes better on a held-out range of population sizes. **(f)** Wasserstein distance between allele frequency spectra at increasing population sizes; convergence occurs by *N ≈* 20,000, which we use for all simulations. **(g)** Example allele frequency spectrum of a simulated sample. **(h)** Scaled allele frequency spectra and interquartile range for PCAWG samples with sufficient number of (> 20) of observed ecDNA-region mutations. The number on the y-axis is the AFS fraction multiplied by the corresponding allele frequency.

A population evolving according to a Wright-Fisher model with constant population size would display the classical *f^−^*^1^ tail in the AFS^18^. In contrast, the tumor population expands exponentially. The passenger mutations generated during early expansion expectedly formed a ‘neutral tail’ in the AFS, decaying as *f^−^*^2^ for *f <* 0.5 (Figure 1b) ^14^. The clonal (somatic, heterozygous) mutations clustered with variant allele frequency (VAF) centered at 0.5 (e.g., Figure 1c).

We next introduced a single emerging subclone with increased fitness at time *T*_1_. We denote its *emergence time* using *g* (= *T*_1_ − *T*_0_), measured in number of population doublings. The additional fitness was parameterized by a *selection strength s*, implying that the subclone was (1 + *s*)× more likely to divide than a cell from the founder tumor clone. The subclone emergence results in a mixed population consisting of the original founder clone and a positively selected lineage descended from the cell in which the subclone occurred (Figure 1a). This event generates a distinct second mode in the allele frequency spectrum (AFS), positioned between the clonal peak and the neutral tail (Figure 1d). The exact location of the subclonal peak is determined by the eventual size of the subclonal population relative to the total population, and therefore depends jointly on *g* and *s* (Figure S1a). The width of the (sub-)clonal peak is governed primarily by binomial sampling variability from finite sequencing depth; higher coverage reduces this sampling noise and sharpens the mode (Figure S1b).

In addition to *g* and *s*, other parameters also shape the AFS. The number of clonal mutations *C* carried by the founder cell directly determines the height of the clonal peak, while mutations accumulating neutrally with rate *µ* (expected number of somatic mutations in a haplotype per division) during tumor growth, contribute to the characteristic *f^−^*^2^ tail (Figure S1c). Both *g* and *µ* determine the height of the subclonal cluster relative to the clonal cluster. Similarly, *g* and *s*determine the size of the subclonal population, and hence the location of the subclonal cluster at *ϕ/*2, where *ϕ* denotes the *descendant fraction* or the fraction of cells descended from the subclone (Figure 1a). The combined effects of emergence time *g*, selection strength *s*, mutation rate *µ*, and initial clonal load *C* therefore determine the patterns observed in the AFS. The width of the AFS peaks is further influenced by *sequencing depth*, and the AFS might itself shift on the x-axis, based on sample *purity*, defined as the fraction of cells that are descendants of the tumor clone in the sample. We exploited these ideas to analyze the behavior of the AFS in the special case where the subclone arises due to ecDNA formation.

### Efficient simulation allows sampling of ecDNA evolution

We used forward simulation to generate data for parameter estimation algorithms. Each simulation takes as input the evolutionary parameters (*s*, *g*, *µ*, *µ_e_*) and produces a pair of 100-bin histograms representing the AFS at each of four sequencing depths (30×, 50×, 100×, 200×), obtained by binomial resampling of the final cell population. Forward simulation of ecDNA dynamics requires explicit tracking of chromosomal mutations, ecDNA-specific mutations, and stochastic ecDNA segregation. Because training a deep learning inference model requires generating over a million simulated tumors, simulations must be sufficiently efficient to make large-scale data generation practical.

The runtime depends not only on the population size *N*, but also on the evolutionary parameters. Higher selection or lower emergence, for example, led to many copies of ecDNAs being created earlier on, which required additional bookkeeping and processing for each cell division. To characterize runtime scaling, we first simulated 20 tumors across a grid of evolutionary parameters and measured time taken for each simulation to reach different population sizes between 0 and 100, 000. With efficient data structures (Methods), the simulations scaled as ∼ *N* log(*N*), allowing us to simulate N=100K cells in 2 mins, and N=20K cells in ∼ 15s(Figure 1e).

We next asked how large the simulated population needed to be for the allele-frequency spectrum (AFS) to stabilize. Using the Wasserstein distance between AFS obtained at increasing population sizes and the spectrum at a final population size of N=50K, we found rapid convergence by N=20K (Figure 1f). Beyond this point, the spectra changed minimally, and the positions of clonal, subclonal, and ecDNA-amplified components were largely unchanged. Based on these results, we fixed the final population size at N=20K cells for all subsequent simulations. Sequencing coverage also shapes the information available for inference: at lower depths, binomial sampling noise broadens the AFS peaks, reducing resolution for both subclonal detection and parameter estimation. Therefore, we simulated AFS using three representative depths (30×, 50×, and 100×) and covering parameter ranges *s* ∈ [0, 2.5], *g* ∈ {0, …, 9}, *µ* ∈ {1, …, 500} mutations per division, and *µ_e_* ∈ [0.01, 3].

### EcDNA amplification produces two distinct and jointly informative AFS

EcDNA is a special type of subclone, because it influences mutations in a large (∼ 2Mbp) region, not just a point location. Its formation is often followed by a burst of clustered mutations (kataegis). Because such events are likely to occur at formation, the resulting mutations are all ancestral to the ecDNA-bearing lineage and therefore appear as early, high-frequency variants. The AFS of mutations on ecDNA are best investigated separately. We refer to the AFS restricted to the ecDNA-overlapping region as *ecAFS* and to the diploid genome spectrum as the *chromosomal AFS (chAFS)*.

Random segregation and selection result in high copy numbers of ecDNA per cell^6^, which can substantially distort the ecAFS relative to the standard diploid expectation. If an ecDNA-bearing lineage carries an average of *K* ecDNA copies per cell, then a mutation present on all (or most) of those copies has expected allele frequency *K/*(*K* + 2) (Figure 1a). For large *K*, this value approaches 1, implying that sequencing reads from the ecDNA-overlapping region would be dominated by ecDNA rather than chromosomal templates. We therefore hypothesized that the ecAFS would carry a prominent cluster of high-frequency mutations arising from ecDNA-amplified alleles. Conversely, mutations that remain confined to the diploid chromosome—such as clonal mutations present on exactly one homolog prior to ecDNA formation—are diluted by the expanding pool of ecDNA copies. Somatic mutation on a single chromosome has expected allele frequency 1*/*(*K* + 2). Therefore, we additionally hypothesized that intermediate-frequency variants in the ecDNA-overlapping region would largely disappear, collapsing toward either very low frequencies—mutations present on the chromosomal copy of the region, or arising on a single ecDNA copy and diluted as the copy number grows—or very high frequencies (mutations carried by most ecDNA copies). Allele frequencies elsewhere in the genome would continue to exhibit the standard mixture of clonal, subclonal, and neutral variants. These predicted signatures were indeed observed in simulations (Figure 1g).

### Empirical AFS from TCGA/PCAWG tumors were consistent with the simulations

The chAFS and ecAFS data observed in simulations matched the corresponding AFS in ecDNA-positive TCGA/PCAWG tumors(Figure 1h). Specifically, the ecAFS displayed both (i) a broad high-frequency component corresponding to ecDNA-amplified mutations and (ii) a marked depletion of intermediate-frequency chromosomal variants prior to ecDNA formation. Tumors lacking ecDNA showed no such distortions. Moreover, the chAFS showed the expected behavior of subclone formation, despite the subclone arising due to ecDNA. The close agreement between the simulated and tumour-mutation derived spectra validated our forward simulation model in capturing the mutation and amplification dynamics characteristic of ecDNA-bearing tumors. The somatic mutations in simulations and in the empirical data were over the same range (Supplementary Figure S3), although more uniformly distributed in simulations, to ensure sufficient training data for all parameter ranges. The average copy number of ecDNA per cell was much lower in simulations relative to TCGA/PCAWG. This is partly because of a simpler fitness parameter, which incentivizes presence of ecDNA but not ecDNA count. Notably, this does not change the AFS shape dramatically. The peak of the high frequency mutations in ecAFS is located at *K/*(*K* + 2). As the number of ecDNA per cell changes, the location changes by a small amount, for example from 0.83 (K=10) to 0.96 (K=50), always well separated from the other somatic mutations, whose frequency is below 0.5.

### Purity-aware data augmentation improves robustness to tumor purity misestimation

Allele frequencies in bulk sequencing represent a mixture of tumor and normal DNA, and the degree of dilution depends on tumor purity. Although PCAWG provides purity-corrected allele fractions, misestimation of purity can distort the allele frequency spectrum in both ecDNA-overlapping and non-ecDNA regions. In particular, overestimation of purity inflates observed allele frequencies, while underestimation compresses them toward zero, systematically altering the shapes of clonal, subclonal, and ecDNA-related components (Supplementary Figure S2).

Because our deep learning models are trained on simulated spectra assuming 100% tumor purity, a direct application to real tumors would introduce a train–test shift. To mitigate this, we explicitly modeled the effect of purity errors during training by randomly perturbing the AFS according to the purity and its mis-estimation (Methods). These transformations were performed in a copy-number–aware manner and applied independently to the ecDNA-overlapping and diploid genomic regions.

The resulting perturbed spectra retained the qualitative evolutionary signals (overall shape) while incorporating the distortions expected (leftward or rightward shift, and/or tightening) (see Supplementary Figure S2) from imperfect purity correction. With the purity augmented simulated spectra as input, we asked if the key parameters of the model (*s, g, µ, ϕ*) could be estimated.

### Cross-attention based neural networks estimate model parameters

We trained models that accept chAFS and ecAFS as inputs— each a 100-bin histogram — together with the ecDNA-region copy number as a scalar feature. To estimate the ecDNA descendant fraction *ϕ̂* prior to inference of other parameters, we trained a separate one-dimensional convolutional neural network called *ecEvol-CNN* (Supplementary Figure S4a and Methods). To estimate other parameters (*ĝ, ŝ, µ̂*), we applied a cross-attention inference model, *ecEvol-CA* (Figure 2a, Supplementary Figure S4b and Methods). The two methods were combined in a pipeline called *ecEvol*. ecEvol accepts chAFS and ecAFS as input and outputs (*ϕ̂, ĝ, ŝ, µ̂*). It works with either simulated input for training, validation, and testing, or input derived using somatic mutation calling on tumour wgs samples (Figure 2a).

**Figure 2:**
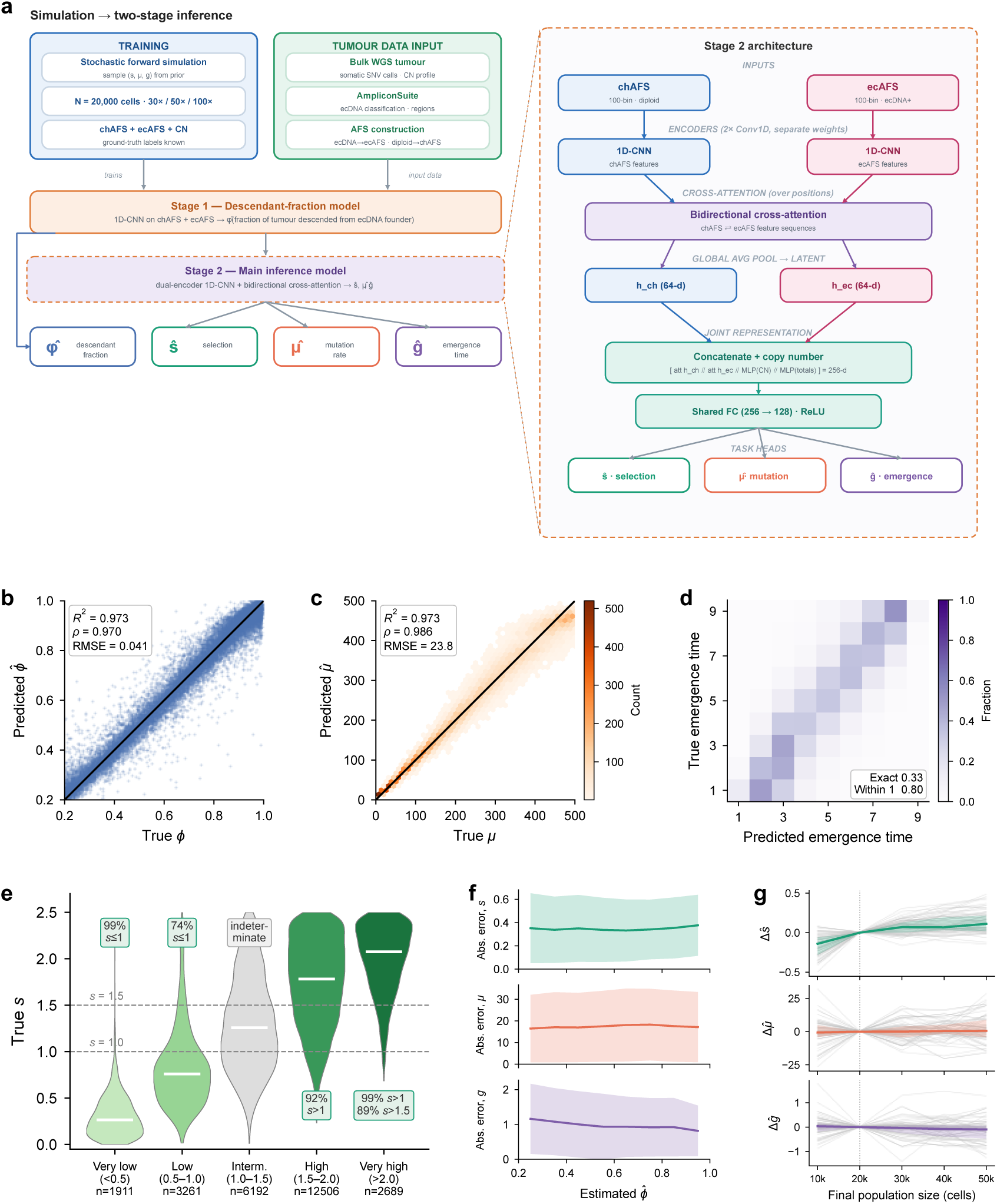
Cross-attention model recovers ecDNA evolutionary parameters. **(a)** Simulation-based training and two-stage inference pipeline. A stochastic forward simulator generates paired chromosomal/ecDNA allele-frequency spectra (chAFS/ecAFS) with known (*s, µ, g*); for real tumours, AmpliconSuite identifies ecDNA(+) features and the spectra are constructed from somatic SNVs. Stage 1 estimates the descendant fraction *ϕ̂*; Stage 2 (dual-encoder 1D-CNN with bidirectional cross-attention over the spectra, combined with copy number) jointly infers *ŝ, µ̂, ĝ*. **(b)** Predicted vs. true descendant fraction *ϕ̂* (*R*^2^ = 0.973, Spearman *ρ* = 0.970, RMSE = 0.041). **(c)** Mutation rate *µ̂* (*R*^2^ = 0.973, *ρ* = 0.986, RMSE = 23.8). **(d)** Confusion matrix for emergence time *ĝ* (exact recovery 33%; within one generation 80%). **(e)** True *s* stratified by predicted confidence tier, showing high positive predictive value at the extremes. **(f)** Absolute prediction error as a function of estimated *ϕ̂*; errors remain stable across the range for all three parameters. **(g)** Change in each inferred parameter when the same simulated tumor is observed at a different final population size, relative to that simulation’s own 20,000-cell estimate (*n* = 70 simulations, each checkpointed at all five sizes). Faint lines are individual simulations, the s2o8lid line the median, the band the interquartile range, and the dotted line the training population size.

Model performance improved monotonically with sequencing depth. At 100× sequencing depth, the final model achieves the strongest recovery of selection strength (*R*^2^ ≈ 0.53, MAE ≈ 0.36), low error for mutation-rate inference (*R*^2^ ≈ 0.97, MAE ≈ 17.3), and strong tolerance for near-miss errors in emergence time (off-by-one accuracy ≈ 0.80). Performance degraded gracefully at lower depths, consistent with increased sampling noise. Unless otherwise stated, results on simulations below are shown for hold-out test data with sequencing depth 100×, which we use as a representative setting; performance across additional depths is reported in Supplementary tables S1, S2, S3.

ecEvol-CNN accurately estimated *ϕ* (*R*^2^=0.97; Figure 2b). Similarly, ecEvol-CA accurately estimated mutation rates *µ* across the full simulated range (*R*^2^=0.97; Figure 2c). Emergence time estimates *ĝ* tracked the true *g* values (Supplementary Figure S5a). When *ĝ* values were rounded to the nearest integer, they matched the correct emergence time exactly in 33% of cases and within one generation in 80% of cases (Figure 2d).

Estimated selection coefficients *ŝ* also tracked the true values, with some dispersion and saturation at high *s* values (Supplementary Figure S5b). Partitioning *ŝ* into five regimes using cutoff values at (0.5, 1.0, 1.5, 2.0), we observed that predictions in the high-selection regimes (*ŝ* ≥ 1.5) corresponded to true *s >* 1 in 93% of cases, while low-selection estimates (*ŝ <* 1) corresponded to true *s <* 1 in 83% of cases (Figure 2e). Intermediate predictions were ambiguous. Notably, while the training was limited to samples with *ϕ <* 0.99, the prediction error remained stable across the range of *ϕ̂* (Figure 2f), showing strong generalization. Predictions were also stable when the same tumor was observed at a different final population size. Rescoring 70 simulated tumors checkpointed at population sizes from 10,000 to 50,000 cells shifted the median estimate by at most 5.7% of a parameter’s range relative to its 20,000-cell value, and in every case by less than the variation between simulations (Figure 2g).

### Ablations justify the dual-spectrum input, cross-attention, and purity-aware training

We ablated the model’s key design choices on held-out simulations at 100× depth (Supplementary Table S4). To establish which differences were meaningful, we first retrained the final architecture with ten random seeds, which gave a standard deviation of 0.004 in Spearman *ρ_s_* (range 0.662 to 0.673). Differences in *ρ_s_* above roughly 0.01 therefore exceed initialization variance.

Supplying both spectra was necessary. Given only the chromosomal spectrum (chAFS), the model achieved *ρ_s_*= 0.641, compared with 0.670 when both chAFS and ecAFS were provided. Allowing the two spectra to interact through cross-attention gave a further improvement over a dual-encoder CNN without attention and over a two-channel CNN (*ρ_s_* = 0.670 versus 0.650 and 0.643). Purity-aware training produced the largest effect. Without augmentation, the model matched the augmented model on clean data (*ρ_s_* = 0.670), but its selection recovery collapsed under purity misestimation (*ρ_s_* = 0.431), whereas the augmented model was essentially unaffected (0.670 to 0.665). Multi-task training matched a single-task *s* model (0.670 versus 0.666, within seed variability) while additionally yielding *µ* and *g* from the same network. The two cross-attention variants were statistically indistinguishable (0.670 versus 0.665), and we selected the pooled model, which achieved the same accuracy with 118,572 parameters compared with 447,468 for the flattened variant.

To benchmark the deep learning approach, we implemented a rejection-based Approximate Bayesian Computation (ABC) baseline using the same simulated reference library (Methods). On held-out test data, the deep learning model outperformed ABC on all three parameters: Spearman *ρ* for *s* was 0.665 (DL) versus 0.603 (ABC), *R*^2^ for *µ* was 0.967 versus 0.931, and off-by-one accuracy for *g* was 0.79 versus 0.70 (Supplementary Table S5). To assess robustness under purity misestimation, we evaluated both methods on a purity-perturbed test set. The deep learning model trained with purity augmentation retained strong performance (Spearman *ρ* = 0.660), whereas a model trained on clean data deteriorated severely (*ρ* = 0.431, 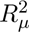 falling from 0.970 to 0.647 and RMSE*_µ_* rising from 25.2 to 86.1), demonstrating that purity-aware training is critical. An ABC variant with a purity-augmented reference library outperformed standard ABC on the perturbed test set (*ρ* = 0.578 versus 0.350), but the purity-augmented deep learning model remained superior under both conditions. These results suggest that in realistic clinical settings where tumor purity is imperfectly estimated, the purity-augmented deep learning model is substantially more reliable than ABC-based inference.

### EcDNA typically emerge early and have high fitness in primary tumors

The Amplicon- Suite pipeline ^19^ has been used to identify ecDNA in many pan-cancer genome projects, including The Cancer Genome Atlas (TCGA), Pan-cancer Analysis of Whole Genomes (PCAWG), Hartwig Medical Foundation (HMF), and other large cohorts^7,12,20^. The PCAWG project is mostly comprised of treatment-naive cases^7^. EcEvol was applied to each of the 385 ecDNA features (288 tumours) to estimate (*ϕ̂, ŝ, µ̂, ĝ*). The median sample returned *ĝ* = 3.67, *ŝ* = 1.91, and *ϕ̂* = 0.94 (Figure 3a), suggesting that the typical ecDNA emerged early, was strongly selected, and expanded to dominate the tumour cell population (Supplementary Data 1).

**Figure 3:**
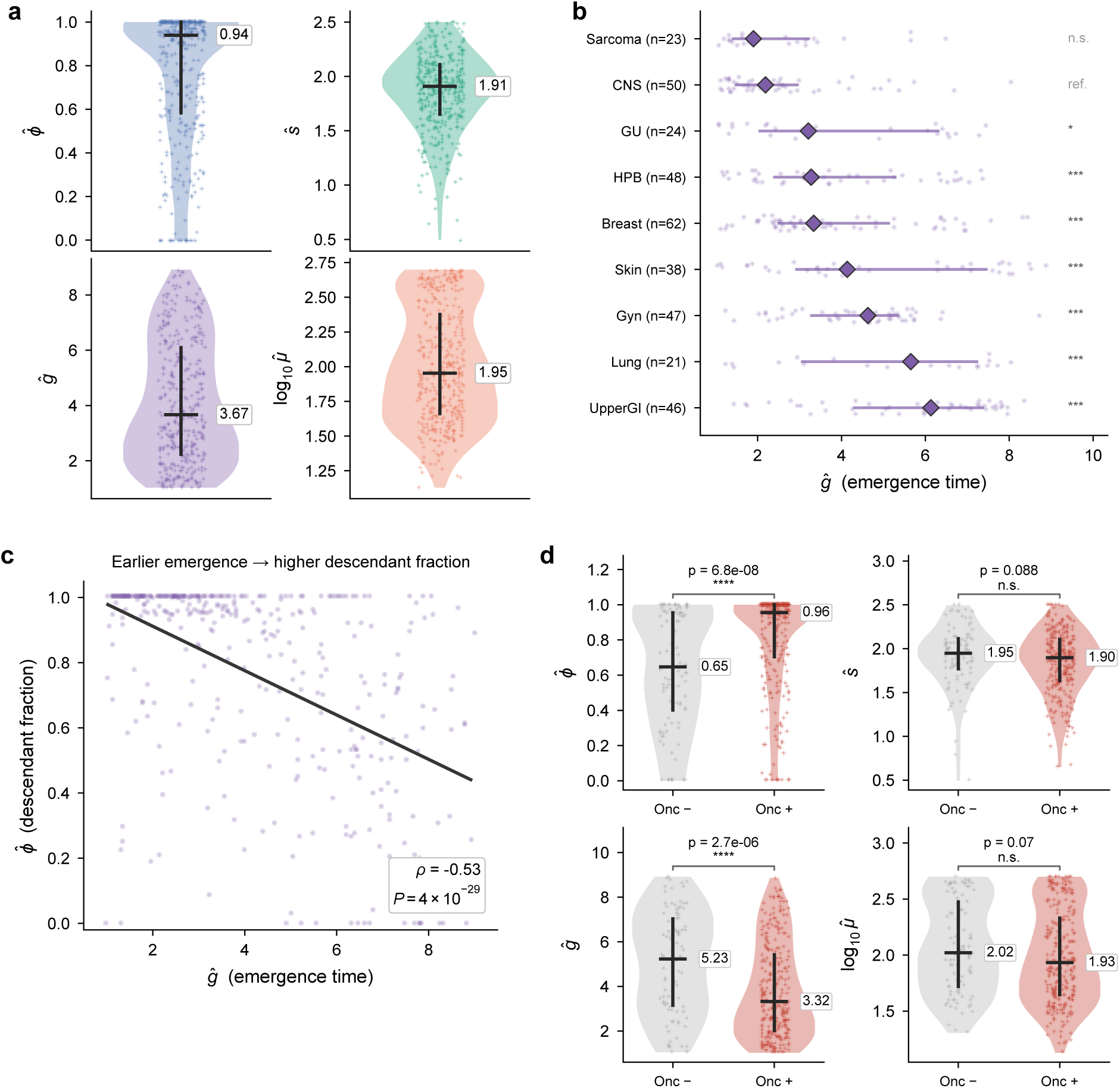
**(a)** Distribution of the four inferred parameters across scored ecDNA features; selection *ŝ* is tightly concentrated (∼1.9) while descendant fraction *ϕ̂*, emergence time *ĝ* and mutation rate *µ̂* vary widely. **(b)** Emergence time *ĝ* by cancer type (diamonds, median; bars, interquartile range; groups with *n* ≥ 15 ecDNA). Emergence time differed between groups (Kruskal–Wallis *P* = 1.4 × 10*^−^*^12^); symbols give Mann–Whitney *U* tests against CNS, Benjamini–Hochberg adjusted (\**P <* 0.05, \*\*\**P <* 0.001; n.s., not significant). EcDNA arose earliest in CNS and sarcoma and latest in lung and upper-gastrointestinal tumours. **(c)** Earlier-emerging ecDNA reach a higher descendant fraction (Spearman *ρ* = −0.53, *P* = 4.4 × 10*^−^*^29^; line, least-squares fit). **(d)** Parameters by ecDNA oncogene status: oncogene-carrying ecDNA reach higher dominance and emerge earlier, while per-carrier selection and mutation rate are unchanged (Mann–Whitney *U*).

The four parameters varied to different degrees. Estimated selection coefficients were distributed across the positively selected regime (interquartile range 1.65–2.11, full range 0.49–2.50), with 85% of samples exceeding *ŝ* = 1.5, while *ϕ̂*, *ĝ* and *µ̂* spanned wider portions of their ranges (Figure 3a). The concentration of *ŝ* at higher values was anticipated from the way ecDNA-positive samples are ascertained. EcDNA is called from bulk sequencing only after the ecDNA-bearing lineage has expanded enough to alter the copy-number and allele-frequency profile, so the cohort was conditioned on high clonal dominance (median *ϕ̂* = 0.94; 55% of samples with *ϕ̂* ≥ 0.9). The same conditioning shifted the selection distribution in simulations, where the generating parameters were known. Although training parameters were drawn uniformly from *s* ∈ [0, 2.5] (median 1.50), simulated tumours that reached *ϕ* ≥ 0.9 had a median true *s* of 1.73. Samples in which ecDNA is detectable are therefore expected to be enriched for stronger selection, and the observed distribution was consistent with this ascertainment.

Estimates also varied across tumour types (Supplementary Figure S6). Median *ŝ* ranged from 1.55 to 2.15 across tumour groups, while median *ĝ* varied more than three-fold over the same groups (Figure 3b). EcDNA emerged earliest in sarcoma and CNS tumours (median *ĝ* of 1.90 and 2.19 doublings) and much later in lung and upper-gastrointestinal tumours (median *ĝ* of 5.65 and 6.13). The early emergence in CNS tumours was consistent with the aggressive and early-onset biology of these cancers, in which ecDNA amplification is acquired as an early event^12^. In these tumours, the model placed most of the posterior probability at the earliest identifiable generation (median *P* (*ĝ* ≤ 2) = 0.66, against 0.33 elsewhere), and in 68% the 90% credible interval reached the earliest value the model can express. Since estimates near the lower limit of *g* are compressed upward (Supplementary Figure S5a and Methods), the true emergence of these ecDNA is likely to be at least as early as estimated, in line with their reported truncal origin. The early emergence in sarcoma was mediated by ecDNA carrying MDM2 or CDK4 (*n* = 16). For these samples, median *ĝ* = 1.64 and *P* (*ĝ* ≤ 2) = 0.78. Moreover, in 81% of cases, the 90% credible intervals reached the earliest value that ecEvol can estimate. Sarcoma ecDNA that did not carry MDM2/CDK4 emerged later (median *ĝ* = 3.13, *n* = 7; *P* = 0.033, Mann–Whitney *U*), as did the remaining PCAWG ecDNA outside sarcoma (median 3.80, *P* = 1.2 × 10*^−^*^4^).

The lung group was predominantly squamous (17 of 21 ecDNA amplicons), with a low descendant fraction suggesting subclonality (median *ϕ̂* = 0.53). Earlier results that timed chromothripsis patterns have also suggested that chromothripsis followed by oncogene amplification were later events in lung squamous cell carcinomas when compared to acral melanomas^21^, consistent with ecEvol. The larger median times in upper-GI cancers is intriguing because ecDNA often arise early, often in a precancerous dysplasia (Barrett’s esophagus) ^8^. Their early occurrence in BE is strongly associated with progression to esophageal cancer. However, the study also noted that ecDNA incidence in esophageal cancer increased from 25% of early-stage to 43% in late-stage tumours. Our results suggest that the newly emerging ecDNA can skew the median emergence time higher.

Emergence time and clonal dominance were related across the cohort. EcDNA that emerged earlier reached higher descendant fractions (*ρ* = −0.53 between *ĝ* and *ϕ̂*; Figure 3c), consistent with early-arising ecDNA having more of the tumour’s growth history over which to expand.

Denote ecDNA-positive samples as ecDNA-onc^+^ (oncogene-positive) if their ecDNA amplify a known oncogene. They can be contrasted with ecDNA-onc*^−^* samples, which are ecDNA-positive but the ecDNAs do not amplify a known oncogene. EcDNA-onc^+^ samples emerged earlier (median *ĝ* = 3.32 versus 5.23, *P* = 2.7 × 10*^−^*^6^) and reached a higher descendant fraction (median *ϕ̂* = 0.96 versus 0.65, *P* = 6.8 × 10*^−^*^8^, Mann-Whitney) than ecDNA-onc*^−^* samples, while the inferred selection coefficient and mutation rate did not differ significantly (Figure 3d).

### Advanced tumours are characterized with later ecDNA emergence

We next analyzed advanced cancers in the HMF cohort (*n* = 1,336 ecDNA features from 913 samples of 858 patients; Supplementary Data 1), and compared estimates to PCAWG. The most striking finding was the later emergence of the median HMF ecDNA (Figure 4a). The overall cohort medians were *ĝ* = 3.67 in PCAWG, but 7.02 in HMF, close to a two-fold increase. Inferred selection coefficients were higher in HMF (median *ŝ* = 2.15 versus 1.91 in PCAWG, *P* = 6.3 × 10*^−^*^63^, Mann-Whitney). Despite the HMF ecDNA having higher fitness, the late emergence resulted in a smaller descendant fraction, with a median *ϕ̂* of 0.76 compared to 0.94 for PCAWG (*P* = 1.6 × 10*^−^*^6^, Mann-Whitney).

**Figure 4:**
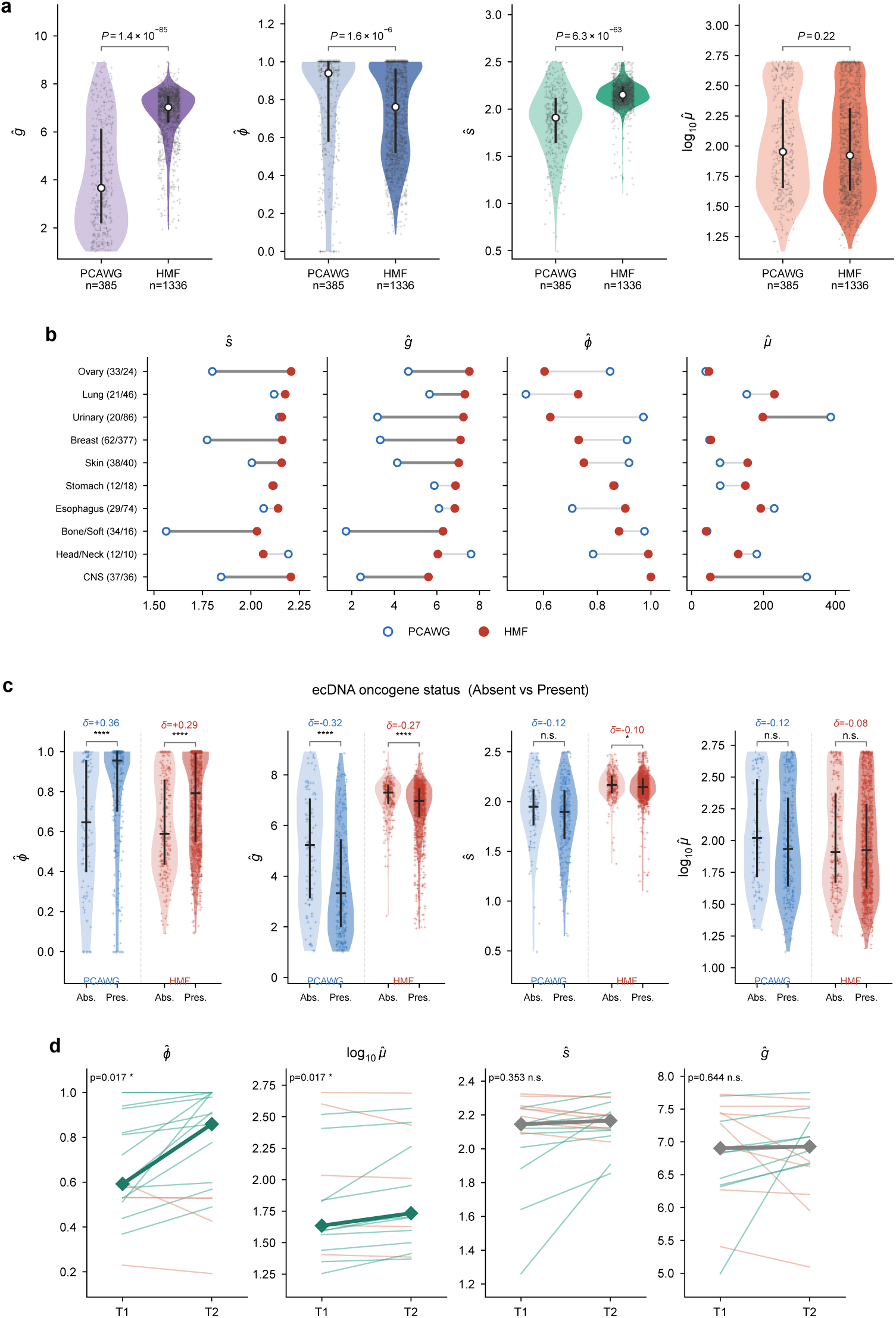
ecDNA evolution in the HMF advanced/treated cohort. **(a)** Distribution of the four inferred parameters across all scored ecDNA features in PCAWG (light; *n* = 385) and HMF (dark; *n* = 1,336). *P* values from two-sided Mann–Whitney *U* tests. **(b)** Per-cancer-type shift from primary (PCAWG, open) to advanced/treated (HMF, filled) for each parameter (matched cancer types; dark connectors indicate FDR *<* 0.05, Mann–Whitney *U* with Benjamini–Hochberg correction). **(c)** Parameters by ecDNA oncogene status in both cohorts (absent vs. present; Cliff’s *δ*). **(d)** Within-patient longitudinal change across timepoints (T1→T2); descendant fraction *ϕ̂* and mutation rate increase while selection and timing are stable (Wilcoxon signed-rank).

To test for confounding differences due to subtypes, we additionally mapped PCAWG tumor types onto the 10 HMF cancer groups, merging codes where a single HMF group corresponded to more than one PCAWG type, and retained only groups with at least five ecDNA features in each dataset (Methods). The HMF group medians for emergence time exceeded their PCAWG counterparts in 9 of the 10 matched groups (Wilcoxon signed-rank, *P* = 9.8 × 10*^−^*^3^; Figure 4b; Supplementary Figure S6). The shift was also evident within single matched disease entities: for example, the CNS groups comprised glioblastoma in both cohorts, and the median emergence time rose from *ĝ* = 2.41 in PCAWG to 5.60 in HMF with non-overlapping interquartile ranges (*P* = 6.3 × 10*^−^*^9^, Mann-Whitney). The only group in which emergence time did not increase was head and neck, which was also among the smallest cohorts (*n* = 12 and 10 samples). For selection, the HMF group medians exceeded PCAWG in 8 of 10 groups (Wilcoxon signed-rank, *P* = 0.027; Figure 4b), but the shift was small relative to the within-group spread, and both cohorts had an ascertainment bias for detectable ecDNA. The shift was not immediately explainable by oncogene presence, as the differences showed up even when comparing samples amplifying the same oncogene (Supplementary Figure S7).

To partially explain these differences, we checked the treatment status of the 733 samples that were ecDNA-positive and had a recorded treatment status. Unlike the treatment-naive PCAWG samples, 627 of the 733 HMF samples were pretreated and the remaining had advanced or metastatic tumors. Kim et al^7^ had previously observed that the ecDNA count per tumor was significantly higher in untreated HMF tumors compared to the primary cancer found in PCAWG, was even higher in pre-treated samples compared to untreated samples and also increased with the number of treatments. Similar observations have been made associating ecDNA formation and evolution with drug treatment^10,22^. Our results support the hypothesis that new ecDNA arising after treatment or upon cancer progression skewed the emergence time higher.

These inter-cohort differences were robust to sample quality: restricting both cohorts to samples sequenced at ≥50× with purity ≥0.60, the purity range used for training augmentation (159 PCAWG and 447 HMF features), reduced but did not eliminate the gaps in *ĝ* (median 3.89 versus 7.01, *P* = 1.3 × 10*^−^*^30^), *ϕ̂* (0.94 versus 0.85, *P* = 0.016) and *ŝ* (1.89 versus 2.17, *P* = 1.1 × 10*^−^*^32^; Supplementary Figure S8). Taken together, the results are consistent with earlier observations that new ecDNA-positive cases arise as the tumour evolves ^7,12^. Our results also suggest the new ecDNA often have higher fitness and rapidly colonize the existing population.

The oncogene comparison in HMF reproduced the pattern seen in PCAWG (Figure 4c). EcDNAonc^+^ samples emerged earlier (median *ĝ*=6.97 for onc^+^ versus 7.30 for onc*^−^*, *P* = 5.5 × 10*^−^*^11^) and reached a higher descendant fraction (median *ϕ̂*=0.79 versus 0.59, *P* = 4.0 × 10*^−^*^13^, Mann-Whitney). In contrast, the inferred selection coefficient differed only marginally (median 2.15 versus 2.17, *P* = 0.017) and the mutation rate did not differ significantly. Thus, the timing and extent of ecDNA expansion in both the primary and the advanced cohorts were associated with the amplification of known oncogenes on the ecDNA.

The HMF data also contains longitudinal samples with the same patient biopsied at multiple time points. We matched ecDNA features across time points within each patient and retained the best-matching pair per patient, yielding *n* = 17 paired samples with a median interval overlap of 0.99. The descendant fraction increased from T1 to T2 (median *ϕ̂* shifted from 0.59 to 0.86; increase in 11 of 17 patients, Wilcoxon signed-rank *P* = 0.017; Figure 4d), consistent with continued expansion of the ecDNA-bearing lineage between biopsies. The inferred mutation rate also increased (median *µ̂* changed from 43 to 53 mutations per division; increase in 12 of 17 patients; *P* = 0.017), although the magnitude of this change was small relative to the range over which *µ̂* was estimated. The median paired increase was 7.3 mutations per division, or 1.5% of the trained range of 1 to 500 mutations per division, against a spread of 18 to 492 across patients at T1. In contrast, the two parameters that describe the origin of the ecDNA rather than its subsequent expansion were unchanged between biopsies: emergence time (median *ĝ*: 6.90 to 6.93, *P* = 0.64) and selection coefficient (median *ŝ*: 2.15 to 2.17, *P* = 0.35) were both stable. Because the same ecDNA was tracked within a patient, *ĝ* and *ŝ* were expected to remain constant, and their stability across independent biopsies provided an internal check on the reproducibility of the estimates.

### Early ecDNA emergence is associated with adverse overall survival

EcDNA-positive tumours had previously been compared against a reference group of PCAWG patients carrying non-ecDNA amplifications^20^ (linear amplification, BFB, or complex non-cyclic structures, but no ecDNA; *n* = 1,448), and showed markedly worse overall survival than this reference (*P <* 0.001), indicating that ecDNA *per se* conferred a survival cost beyond focal amplification. We next asked whether estimated ecDNA evolutionary parameters imposed a differential survival burden.

We restricted the analysis to patients with a single scored ecDNA (*n* = 170, 85 events). Tumours whose ecDNA emerged earliest (lowest *ĝ* tertile) showed worse survival than the rest (60% vs 45% events; log-rank *P* = 3.3 × 10*^−^*^4^; Figure 5a). The association held after stratifying by cancer type and adjusting for age, sex, and copy number (Cox HR = 1.87, 95% CI 1.11–3.14, *P* = 0.018), but weakened when CNS tumours were removed (*n* = 140, HR = 1.79, *P* = 0.057).

**Figure 5:**
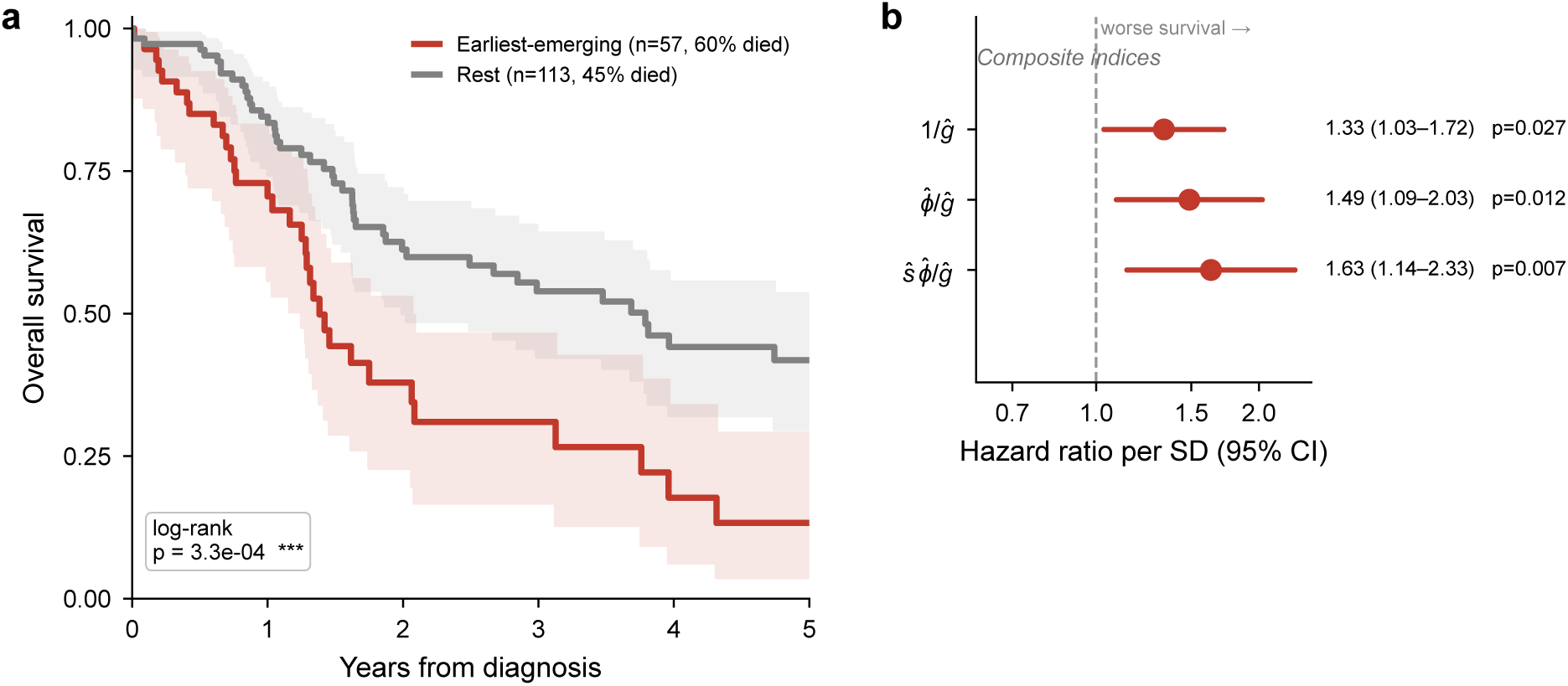
ecDNA emergence timing and overall survival (PCAWG). **(a)** Kaplan–Meier overall survival for tumours in the earliest-emergence tertile vs. the rest (single-scored-ecDNA cohort, *n* = 170, 85 events; log-rank *p* = 3.3 × 10*^−^*^4^). **(b)** Cox proportional-hazards forest for composite evolutionary-aggressiveness indices (1*/ĝ*, *ϕ̂/ĝ*, *ŝϕ̂/ĝ*), stratified by cancer type and adjusted for age, sex and copy number. Nested likelihood-ratio tests show the prognostic signal is carried by emergence timing: adding dominance (*ϕ̂*, *p* = 0.61) or selection (*ŝ*, *p* = 0.72) contributes no further information.

We investigated whether inferred dominance or selection added to this timing signal. In nested likelihood-ratio tests, neither *ϕ̂* (*P* = 0.61) nor *ŝ* (*P* = 0.72) improved upon the association with emergence time alone. However, composite indices of 1*/ĝ*, *ϕ̂/ĝ*, and *ŝϕ̂/ĝ* each stratified survival significantly (Figure 5b). Our results suggest that these indices can be used to order and triage ecDNA(+) patients for tumour pathology.

## Discussion

The presence of ecDNA in tumor cells is a marker of pathology, associated with increased resistance, and worse survival outcomes for patients^7,9,12,20^. However, these published results also point to a high level of variability between ecDNA-positive samples, even when the samples are drawn from the same cancer subtype. All ecDNA are not alike. They may emerge at different times in tumor evolution, may provide differential fitness advantage to the tumor, and may have contributed different numbers of descendant cells to the tumor population at sampling time. Some natural questions arise: Can these evolution parameters be estimated from genomic DNA sampled from an ecDNA-positive tumor? Do the estimated parameters provide insight into the pathology of a specific sample. Our answer to both questions is a qualified yes.

Our manuscript outlines a method, ecEvol, that estimates these key evolutionary parameters of ecDNA evolution. It utilizes two distinct variant allele frequency spectra; a sparse one from the genomic regions comprising the ecDNA which contributes ultra-high and ultra-low frequency mutations, and a separate AFS derived from the rest of the genome, which contributes intermediate frequency clonal and subclonal mutations. We also utilize a novel cross-attention neural network model that learns the dependencies between the different frequency bins both within the same AFS and between the two AFS. In simulations, ecEvol predicts emergence time and descendant-fraction with high accuracy over a wide spectrum of model parameters. It also predicts selection strength to accurately distinguish high-fitness from lower-fitness samples. EcEvol’s performance improves upon all other competing models that we tried.

We recognize that ecEvol utilizes a simple evolutionary model. For example, the selection parameter is constant and depends only on the presence/absence of ecDNA in the cell. A more realistic model would consider the fitness of a cell as a function of the number of ecDNA in it, possibly increasing initially, but subsequently saturating or declining as the number of ecDNA increased beyond what a cell could maintain^6^. With improvements in experimental models of ecDNA evolution ^23^, accurate measurements of ecDNA fitness may be possible. Future work will work with improved fitness functions. We also assume that a single ecDNA arose in the evolutionary history being considered. While most ecDNA positive samples have one ecDNA, others have more, each with its own fitness. Two sub-populations, each with a different ecDNA, but with comparable fitness, would compete, and display confounding AFS signals. Due to the combinatorial expansion of parameters with interacting ecDNA, we chose to work with the simpler model. Even if multiple ecDNA occur, we assume that a single ecDNA with higher fitness and/or early emergence dominates the AFS signal. In future experiments, we will revisit and refine these model choices. In the meantime, the estimated parameters are best interpreted as ‘effective selection strength,’ effective emergence time, and so on. This is analogous to the concept of the notion of effective population size, whose estimates are useful in demographic predictions despite departure from real scenarios.

The tissue of origin and/or cell state, as well as the mutational status of genes encoded within the ecDNA, is likely to play an important role in when an ecDNA arises during the process of cancer. In glioblastoma, in which ecEvol detects early presence of ecDNA, other studies demonstrate that initially ecDNAs arise very early in tumor development as wild type EGFR becomes amplified on ecDNA, but the rapid growth of the tumor seems to correspond to the acquisition of gain of function EGFR mutations occurring in those amplified ecDNAs. In upper GI cancers, for which ecEvol detects a wider spread of time points at which it can arise, including later development of tumors. Importantly, studies in tumor tissue from patients with Barrett’s esophagus sampled over space and time show that ecDNAs can arise during high grade dysplasia in patients who go on to develop esophageal adenocarcinoma, but the initial appearance of the ecDNAs may not yet be transformative ^19^. The development of additional ecDNAs bearing more potent oncogenes may be required for transformation. Other studies have observed that the incidence of ecDNA increased in patients with advanced tumors^7,8,12^. Taken together, these data raise the possibility that ecDNAs may arise at multiple points in tumor development, and the fraction of the cells that contain them depends not only on when they arise, but what is encoded on them, including as new more potent gain of function mutations accumulate on ecDNA particles.

This claim is further bolstered by observations in the HMF cohort, where the majority of samples were obtained from pretreated patients and most of the remaining were advanced, metastatic cases. The incidence of ecDNA increases not only with progression ^8^, but also with treatments^7,12^. Consistent with these observations, our results indeed show a significantly higher emergence time for samples in the HMF cohort relative to PCAWG, and the differences were maintained in matched subtypes. Additionally, we observed good internal consistency in our results in the HMF data. When the same ecDNA in an individual was sampled at multiple time points, we did not see a change in estimated emergence time, but the later time point showed an increased descendant fraction. The later emergence in HMF was maintained even after restricting to specific oncogenes, for nearly all oncogenes tested, suggesting that oncogene presence did not mediate the difference between HMF and PCAWG.

The descendant fraction of ecDNA in the current population is mediated by both, strong selection, and early emergence time. Our results suggested an association between emergence time and survival, and a milder association between selection strength and survival. It is possible that new ecDNA that arose later did not have time to colonize the tumour population, and those tumours were more sensitive to treatment. Overall, these results suggest the prognostic value of modeling ecDNA evolution. As a matter of caution, the survival results were obtained with sparse data and with many covariates. Future studies with larger ecDNA-positive populations should enhance the clinical importance of these findings. In the meantime, ecEvol provides a novel method to understanding the evolution and aggressiveness of ecDNA-positive cancers.

## Online methods

### Forward simulation of ecDNA tumor evolution

We developed a forward-time simulation to model tumor evolution with explicit tracking of chromosomal mutations, ecDNA-specific mutations, and stochastic ecDNA segregation. The simulation begins from a single diploid founder cell and proceeds through repeated birth events until the population reaches a fixed size *N* . Each cell stores the mutations carried on (i) the two chromosomal homologs of the ecDNA-overlapping region, (ii) the two homologs of the non-ecDNA region, and (iii) each individual ecDNA copy after ecDNA formation.

### Cell division and fitness

Each cell divides with a birth rate determined by its ecDNA status. Cells without ecDNA divide at rate *b*_0_ = 1. Cells that carry at least one ecDNA copy divide at rate *b*_1_ = 1 + *s*, where *s* is the selection coefficient. At each simulation step we draw a candidate cell uniformly at random from the current population and accept the division using rejection sampling against the maximum rate 1 + *s*. The candidate divides if

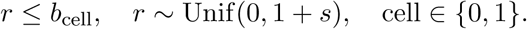

This procedure ensures that ecDNA-positive cells divide in proportion to their fitness advantage.

### Mutation model

Before ecDNA formation, mutations accumulate independently on each chromosomal homolog of both genomic regions as follows: After each cell division, each daughter receives *M*_chr_ new mutations, where

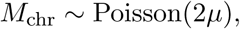

and *µ* is the expected number of mutations per haplotype per mitosis for chromosomal DNA. New mutation identifiers are assigned sequentially.

Once ecDNA is introduced, each ecDNA copy replicates prior to segregation. For a cell with *K* ecDNAs, this produces 2*K* ecDNA copies. Each ecDNA copy accumulates new mutations independently:

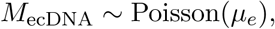

where *µ_e_*is the ecDNA-specific mutation rate. Mutations remain copy-specific and are inherited depending on segregation.

### EcDNA formation and segregation

The parameter *g* is the designated *emergence time* if ecDNA is introduced into a single randomly chosen cell when the population size first reaches 2*^g^*. The ecDNA is created by excising one of the two chromosomal homologs of the ecDNA-overlapping region and assigning it to an initial single-copy ecDNA molecule. The other homolog remains chromosomal.

During each subsequent division, all ecDNA copies are pooled and randomly allocated to the two daughters after replication. Each of the 2*K* ecDNA copies independently selects a daughter with probability 1*/*2, resulting in daughter cells with *K*_1_*, K*_2_ ecDNA, where

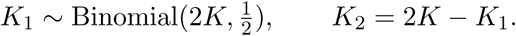

This mechanism permits both stochastic loss and amplification of ecDNA copy number across generations.

### Stopping condition and recorded outputs

The simulation proceeds until the population reaches size *N* . For each mutation, we compute the number of DNA segments on which it appears. In the ecDNA-overlapping region the total number of segments is

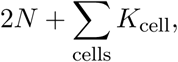

and allele frequencies are defined relative to this quantity. For the non-ecDNA region, the denominator is 2*N* . These frequencies are used to compute the allele frequency spectra prior to sequencing noise and purity distortion.

Allele frequencies are computed in a single pass over all cells at the end of simulation, once the population has reached size *N*, rather than maintaining a global frequency table during growth. Each ecDNA is represented as an individual object carrying its own mutation list, so that copy-specific mutations are faithfully inherited through stochastic segregation. Thus, each cell carries multiple mutation lists, one for the chromosome and one for each constituent ecDNA. A cell division is accompanied by the following local operations: (a) new mutations are added to each of the daughter cell chromosomal DNA with rate *µ*. (b) The ecDNA objects are duplicated and segregated among daughter cells. Each ecDNA object acquires additional mutations with rate *µ*_e_. No population wide updates are needed, resulting in a more efficent running time, as reported in the Results section.

### Sequencing model and construction of allele frequency spectra

After the forward simulation completes, we convert true mutation frequencies into observed sequencing measurements. For each mutation we record its true allele fraction *f*, defined by the number of DNA segments carrying the mutation divided by the total number of DNA segments in the corresponding region. In the ecDNA-overlapping region the denominator is

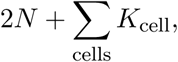

and in the non-ecDNA region it is 2*N* .

For each mutation, we first draw the number of reads covering the locus using a binomial sampling step:

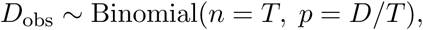

where *T* is the total number of DNA segments in the region and *D* is the target mean depth (30×, 50×, 100×, or 200×). Because *T* is large, this draw is close to a draw with mean *D*. This produces mutation-specific observed coverage values, reflecting the variance expected under binomial sampling of reads from a uniform pool of fragments.

Given the observed coverage *D*_obs_ and the true allele fraction *f*, we draw the number of variant-supporting reads as

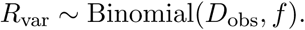

The observed allele fraction is then

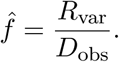

We apply simple quality filters and retain only mutations with *R*_var_ *>* 5 and *f̂ >* 0.01, consistent with typical thresholds used in bulk sequencing-based mutation calling pipelines.

For both the ecDNA-overlapping and non-ecDNA regions, we convert the retained observed allele fractions into 100-bin histograms over [0, 1]. These histograms serve as the inputs to the inference models and are constructed independently for each sequencing depth condition.

We generated a training corpus of 1.26 million such simulated tumors, drawing parameters independently for each run from uniform priors: *s* ∈ [0, 2.5], *g* ∈ {0, …, 9}, *µ_e_* ∈ [0.01, 3], and *µ* ∈ {1, …, 500} mutations per division, with a fixed final population size of *N* = 20,000 cells. Because sequencing noise is applied post hoc to the same final cell population, each forward simulation yields training examples at all four depth conditions simultaneously. To test the impact of population size, we additionally simulated 70 tumors, each checkpointed at final population sizes of 10,000, 20,000, 30,000, 40,000 and 50,000 cells, so that the same tumor is observed at every size.

### Tumor purity perturbation

All simulations are generated under the assumption of 100% tumor purity, and allele fractions represent true frequencies within the tumor. To account for the distortions introduced by purity misestimation in real bulk sequencing data, we apply a two-step perturbation procedure to each simulated allele frequency spectrum.

For each simulation, we draw a “true” tumor purity value

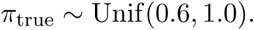

For a mutation with (tumor-only) allele fraction *f*_true_ and tumor-region copy number *C*_tumor_, the expected bulk-observed VAF is

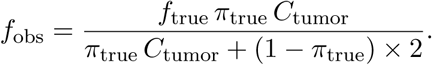

Next, to emulate misestimation, we draw a perturbed purity value

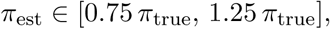

and map the bulk-observed VAFs back into incorrectly corrected allele fractions. The corrected frequency under the wrong purity is

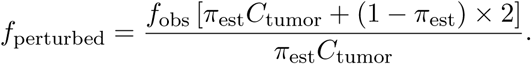

This step is applied independently to both the ecDNA-overlapping region and the non-ecDNA region, using region-specific tumor copy numbers.

After purity perturbation, we recompute the 100-bin allele frequency spectra for each region in the same manner as described above.

### Approximate Bayesian computation baseline

As a non–deep learning baseline, we implemented a rejection-based Approximate Bayesian Computation (ABC) method using the simulated spectra as a reference library. Each library element consists of the concatenated allele frequency spectra from the ecDNA-overlapping and non-ecDNA regions together with the log_2_-transformed ecDNA-region copy number, yielding a 201-dimensional summary vector. Features were standardised to zero mean and unit variance using statistics computed on the training set.

For each test tumor, we compute the same 201-dimensional summary and measure Euclidean distances to all reference summaries in the library. We retain the closest *k* = ⌊*q* · *N*_ref_ ⌋ reference simulations, where *N*_ref_ is the size of the reference library. The tolerance parameter *q* was selected by cross-validation on 1,000 held-out validation simulations, maximising a composite score equal to the sum of the Spearman rank correlation for *s*, *R*^2^ for *µ*, and off-by-one accuracy for *g*, over the grid *q* ∈ {0.0001, 0.0002, 0.0005, 0.001, 0.002, 0.005, 0.01}.

Point estimates are obtained from the accepted set. The selection coefficient *ŝ* and mutation rate *µ̂* are estimated as the arithmetic means of the accepted values; the emergence time *ĝ* is estimated as the mode.

To assess robustness to purity misestimation, we additionally constructed an augmented reference library by appending five purity-perturbed copies of each training simulation using the perturbation procedure described below (“Tumor purity perturbation.”). This expanded the reference library sixfold. The tolerance *q* was re-optimised independently for the augmented reference using the same cross-validation procedure.

### ecEvol: Deep learning inference from allele frequency spectra

We frame inference of ecDNA evolutionary parameters from allele-frequency spectra as a supervised learning problem using simulated tumors with known ground-truth parameters (Fig. 2a). All neural networks are trained exclusively on simulated data and are applied to real tumors only after identical preprocessing and filtering steps.

Because allele-frequency spectra depend strongly on sequencing depth through binomial sampling noise, all models are trained *separately* for each sequencing depth (30×, 50×, and 100×). No parameters are shared across depths.

Two neural network models are trained and applied independently: (i) a 1D convolutional regression model, ecEvol-CNN to estimate the ecDNA descendant fraction *ϕ̂*, reported as a summary of clonal dominance, and (ii) a cross-attention multi-task inference model, ecEvol-CA, to recover ecDNA evolutionary parameters (*ŝ, µ̂, ĝ*), applied to all samples.

Chromosomal and ecDNA allele-frequency spectra are represented as 100-bin histograms over the interval [0, 1]. For the descendant-fraction regression model, spectra are normalized to sum to one so that the network is sensitive only to spectral shape. For the main inference model, spectra are provided without explicit normalization, allowing the network to exploit both shape and amplitude information, which is informative for mutation-rate inference and emergence time.

Training data are augmented to account for tumor purity misestimation as described in the section ‘Tumor purity perturbation’.

### ecEvol-CNN: a descendant-fraction regression model

#### Motivation

The fraction of tumor cells descended from the initial ecDNA-bearing founder, *ϕ*, is a biologically meaningful summary of ecDNA clonal dominance, distinct from selection strength or emergence time. We train a dedicated regression model to estimate *ϕ̂* for each sample so that it can be reported alongside the inferred evolutionary parameters as an independent characterisation of clonal expansion.

#### Architecture

The descendant-fraction model is a one-dimensional convolutional neural network (CNN). The chromosomal and ecDNA allele-frequency spectra are each normalised to sum to one and stacked as a two-channel input of shape (2 × 100). Three Conv1d layers (channels 2 → 16 → 32 → 64; kernel sizes 5, 5, 3; padding 2, 2, 1) with GELU activations and batch normalisation extract spectral features. Adaptive average pooling collapses the frequency axis to a 64-dimensional embedding, which is concatenated with the ecDNA-region copy number scalar to form a 65-dimensional vector. This is passed through two hidden fully connected layers (dimensions 64, 64; GELU activations; dropout rate 0.1) followed by a linear output layer, producing an unbounded scalar. Predictions are clipped to [0, 1] at evaluation time to obtain *ϕ̂*.

#### Training

The model is trained using mean squared error loss against the scaled target *ϕ* = *N*_desc_*/N*, where *N*_desc_ is the number of ecDNA-descended cells at the end of simulation and *N* = 20,000 is the fixed population size. Optimization uses AdamW^24^ (learning rate 10*^−^*^3^, weight decay 10*^−^*^4^) with gradient clipping at norm 1.0 and a reduce-on-plateau learning rate schedule

Training runs for at most 50 epochs and is monitored by the coefficient of determination *R*^2^ on the validation split. The learning rate is halved after three epochs without improvement in validation *R*^2^, training stops after six, and the checkpoint from the best-scoring epoch is retained.

The hyperparameters of this model are listed in Supplementary Table S7. Separate models are trained for sequencing depths 30×, 50×, and 100× using disjoint training, validation, and test splits.

#### Application to real tumors

The descendant-fraction model is applied to all samples to obtain *ϕ̂*. The main inference model is then applied independently to all samples to obtain (*ŝ, µ̂, ĝ*), regardless of the estimated descendant fraction.

### ecEvol-CA. A cross-attention inference model for ecDNA evolutionary parameters

#### Targets

The main inference model predicts three evolutionary parameters: the selection coefficient *s* of ecDNA-bearing cells, the per-division mutation rate *µ*, and the ecDNA emergence time *g*.

#### Architecture

The model employs a dual-encoder architecture with explicit interaction between the chromosomal and ecDNA spectra (Supplementary Fig. S4). Separate convolutional encoders, each two blocks of convolution, batch normalization, ReLU and max pooling, process the two spectra and produce 64-dimensional feature sequences. These are combined using bidirectional cross-attention with four heads, each spectrum attending over the other, to allow the network to learn relative distortions between chromosomal and ecDNA mutation patterns.

Attention outputs are averaged over positions to give one 64-dimensional latent vector per spectrum. The ecDNA-region copy number, log_2_-transformed, and the mutation totals of the two spectra (the count in each region, their sum, and the logarithms of the three) are each embedded to 64 dimensions by two-layer perceptrons. The four components are concatenated into a 256-dimensional joint representation, reduced to 128 dimensions by a shared fully connected layer, and passed to three linear task-specific heads. The network is trained end to end against the multi-task loss below, including the attention weights; no component is pretrained or optimized separately.

#### Outputs and losses

The selection coefficient is predicted on a logistic-transformed scale. With *s*_max_ = 2.5 and *δ* = 10*^−^*^3^, the training target is

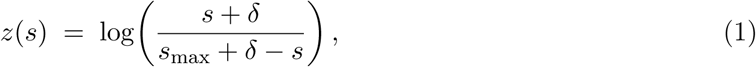

mapping *s* ∈ [0*, s*_max_] to the real line, and the prediction *ẑ* is trained with a Huber loss

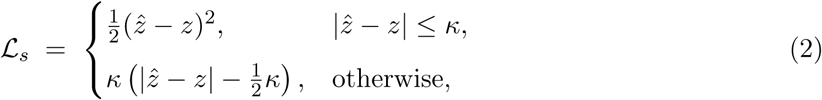

with *κ* = 1. The mutation rate is predicted through a sigmoid output against a min–max-scaled target and trained with mean-squared-error loss, 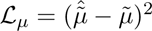. The emergence time is predicted as a softmax over the nine generation classes {0, …, 9} and trained with cross-entropy loss, 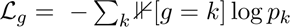, where *p_k_* is the predicted probability of class *k*. The total training loss is the unweighted sum L = L*_s_* + L*_µ_* + L*_g_*, fixed across all experiments.

The emergence-time head outputs a probability over the nine generation classes, and we report *ĝ* as the mean of this distribution rather than the most probable class, which preserves the ordering of samples that share a modal class. The same distribution provides a per-sample uncertainty, and on held-out simulations these probabilities were conservatively calibrated: the smallest set of generations carrying 90% of the probability contained the true emergence time in 94% of cases. Across PCAWG this set spanned three generations for a typical sample. Because the support is bounded, estimates are compressed toward the centre of the range: on held-out simulations the mean *ĝ* was 2.36 for a true *g* of 1 and 7.44 for a true *g* of 9, with negligible bias near the middle. Estimates of early emergence are therefore conservative with respect to the true emergence time.

#### Training procedure

The main inference model is trained using the AdamW optimizer ^24^ (learning rate 10*^−^*^3^, weight decay 10*^−^*^3^, batch size 64, selected as described below) for at most 50 epochs, with gradient clipping at norm 1.0. Training is monitored using the composite validation score defined in Section ‘Model selection and ablation experiments.’ The learning rate is halved after three epochs without improvement in this score, training stops after six, and the best-scoring checkpoint is retained. Separate models are trained for each sequencing depth using fixed random seeds for data splitting and initialization. Full settings are listed in Supplementary Table S7.

#### Model selection and ablation experiments

We evaluated multiple neural network variants on held-out simulated tumors to assess the impact of architectural choices, output parameterizations, and task formulations. All models were trained and evaluated at matched sequencing depth using identical data splits.

A shared two-channel convolutional model serves as a baseline. The selection coefficient was regressed on the logistic-transformed scale with a Huber loss; direct regression on *s* gave negligible differences in accuracy. Similarly, categorical and ordinal formulations for predicting the ecDNA emergence time achieved comparable performance.

Introducing explicit cross-attention between chromosomal and ecDNA representations consistently improved inference accuracy across tasks, including higher rank correlation for selection strength, reduced mutation-rate error, and improved emergence time prediction accuracy.

Removing purity augmentation left clean-data accuracy essentially unchanged but caused selection recovery to collapse on purity-perturbed test data (Spearman *ρ_s_* from 0.67 to 0.43), confirming that augmentation is essential for robustness to purity misestimation.

A single-task model trained to predict only selection strength did not outperform the corresponding head of the multi-task model. Joint training consistently improved stability and accuracy.

#### Model and hyperparameter selection

Architectures and hyperparameters were selected on a held-out validation split using a composite score: the sum of the Spearman correlation and *R*^2^ of *s*, the *R*^2^ of *µ*, and the off-by-one accuracy of *g* (the same criterion used to select the ABC tolerance). On the selected architecture we tuned the learning rate (5 × 10*^−^*^4^, 10*^−^*^3^), weight decay (0, 10*^−^*^4^, 10*^−^*^3^), and batch size (64, 128) over the resulting 12-configuration grid; held-out performance varied by less than 0.01 across the grid. Among architectures that were statistically indistinguishable on the validation composite we selected the most parameter-efficient model, a pooled cross-attention model with ∼119,000 parameters, which we adopt as the final architecture for all depths.

#### Ensemble prediction and reproducibility across random seeds

Individual networks trained from different random initializations reach comparable accuracy but do not assign identical estimates to individual samples. To quantify this, we train ten networks that have identical architecture, hyperparameters, and training data, but differ in random seed, weight initialization and batch ordering. Moreover, the ten models are trained separately for each of 3 sequencing depths (30×, 50×, and 100×), for a total of 30 models. For experimental tumor data, first a depth is chosen (from 30, 50, and 100), that most closely matches the empirical data. The ten corresponding networks for the chosen depth are utilized for estimation. The final estimates provided are the mean across these ten networks.

Ensembling has only a marginal effect on accuracy. On held-out simulations at 100× depth, single networks achieve a Spearman correlation for *s* of 0.669 ± 0.004 (mean ± s.d. over ten seeds), while the ensemble mean achieves 0.676. The corresponding gains are 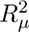 from 0.972 ± 0.001 to 0.975, and off-by-one accuracy for *g* from 0.797 ± 0.005 to 0.804. The tight spread across seeds also provides the reference scale used to interpret the ablation experiments, where differences in *ρ_s_* above roughly 0.01 exceed initialization variance.

The effect on reproducibility is substantially larger. We quantify agreement between networks using the intraclass correlation coefficient (ICC) computed across the ten seeds for every scored ecDNA feature, reporting both the reliability of a single network and, by the Spearman-Brown relation, that of the ten-network mean (Supplementary Table S6). A single network gives an ICC of 0.66 for *s*, 0.75 for *g*, and 0.87 for *µ*, while the ten-network mean gives 0.95, 0.97, and 0.99. Per-sample estimates of *s* from a single network are therefore sensitive to the random seed, and the ensemble is used so that cohort-level analyses do not depend on an arbitrary choice of seed. The descendant fraction *ϕ̂* is estimated by a single Stage 1 network and is identical across the ensemble.

### Inference pipeline for clinical tumor cohorts

#### Cohorts

We applied the inference framework to two whole-genome sequencing cohorts. From the ICGC/TCGA Pan-Cancer Analysis of Whole Genomes dataset we scored 385 ecDNA features across 288 tumors, and from the Hartwig Medical Foundation pan-cancer study we scored 1,336 ecDNA features across 913 tumors. In both cohorts, extrachromosomal DNA was identified using AmpliconSuite-pipeline v0.1344.1, and amplicons classified as cyclic were retained for analysis. PCAWG amplicons were classified with AmpliconClassifier v1.5.1 and HMF amplicons with v0.4.12; both cohorts were processed against GRCh37 with a copy-number gain threshold of 4.5 and a minimum seed size of 50 kbp.

#### Feature construction

For each sample, somatic single-nucleotide variant allele frequencies were partitioned into two sets: those falling within the ecDNA-overlapping genomic region (used to construct the ecAFS) and those falling outside it (used to construct the chAFS). Each spectrum was binned into a 100-bin histogram over [0, 1]. The ecDNA-region median copy number was obtained from AmpliconSuite feature annotations. Samples with fewer than 10 ecDNA-region somatic mutations were excluded prior to inference.

#### Model application

Each sample was matched to the nearest available depth-specific model based on estimated sequencing coverage. The descendant-fraction CNN was applied to obtain *ϕ̂*. The main inference model was then applied independently to obtain (*ŝ, µ̂, ĝ*). Both models were applied to all samples regardless of *ϕ̂*; samples with *ϕ̂* ≥ 0.99 are noted as near-saturated in the results.

### Cross-cohort comparison of inferred parameters

For cross-cohort comparisons, PCAWG tumour types and HMF cancer groups were matched using ten predefined pairings, merging PCAWG codes where one HMF group spans several: CNS (HMF) ↔ GBM (PCAWG), Head and neck ↔ HNSC, Bone/Soft tissue ↔ SARC and BOCA, Esophagus ↔ ESAD, Skin ↔ SKCM and MELA, Stomach ↔ STAD, Breast ↔ BRCA, Urinary tract ↔ BLCA, Lung ↔ LUSC and LUAD, and Ovary ↔ OV.

### Survival analysis

We analysed overall survival in the PCAWG primary-tumor cohort. Deceased patients were assigned an event at their reported survival time, and living patients were censored at the last follow-up interval. Patients missing vital status or a positive time were excluded, as were patients without a recorded age at diagnosis; no values were imputed. The 385 scored ecDNA features correspond to 288 patients, of whom 217 carried exactly one scored ecDNA. We used these single-ecDNA patients, so that no representative amplicon had to be chosen for patients with several ecDNA. After removing patients with missing vital status (*n* = 11), time (*n* = 23), or age (*n* = 13, all sarcomas), 170 patients with 85 events remained.

We split patients at the lower tertile of *ĝ* and compared the earliest-emerging group against the rest by Kaplan–Meier estimation with a log-rank test. The same contrast was then fitted in Cox models adjusted for age, sex, and log_2_ ecDNA-region copy number (age and copy number standardised), with baseline hazards stratified over ten organ-system groups formed from the 27 PCAWG tumor codes; stratifying on tumor code directly would produce several single-patient strata. We repeated this model excluding CNS tumors. The composite indices 1*/ĝ*, *ϕ̂/ĝ*, and *ŝϕ̂/ĝ* were standardised and each fitted in the same adjusted model, and nested likelihood-ratio tests were used to ask whether *ϕ̂* or *ŝ* added prognostic information beyond emergence time.

### Longitudinal analysis of HMF timepoints

A subset of HMF patients were biopsied at multiple time points. For each such patient we compared the earliest and latest biopsy, matching ecDNA features between the two by the Jaccard overlap of their genomic intervals (intersection over union of interval lengths, summed over shared chromosomes). Candidate pairs with overlap below 0.1 were discarded, and the remaining pairs were assigned greedily in decreasing order of overlap so that each feature was matched at most once. Patients carrying several ecDNA yielded several matched pairs; for these we retained only the pair with the highest overlap, so that each patient contributes a single paired observation. This gave 17 patients, with matched pairs overlapping by a median Jaccard of 0.99. Changes between time points were assessed with the Wilcoxon signed-rank test.

## Data availability

Whole-genome sequencing data from TCGA were accessed through dbGaP under accession phs000178. PCAWG/ICGC whole-genome sequencing data are available under controlled access through the ICGC Data Access Compliance Office. Whole-genome sequencing and associated clinical data from the Hartwig Medical Foundation (HMF) were accessed under a licence agreement (HMF DR-057 v3.0); access can be obtained by completing a data request form, available with the application procedure at https://www.hartwigmedicalfoundation.nl/applying-for-data/. AmpliconSuite outputs for PCAWG are available at https://ampliconrepository.org/project/655c060abba7c925095555da. Simulated spectra, train/validation/test splits and trained model weights are available at Mendeley Data (https://doi.org/10.17632/cvcbsvbrn5.1). Per-feature estimates for both cohorts are provided in Supplementary Data 1.

## Code Availability

The ecEvol source code, including the forward simulator, model training and inference, and the scripts that generate all figures and tables, is available at https://github.com/bdamerac/ecEvol. The version used in this study is v1.0.

---

**Supplementary Figure S1:**
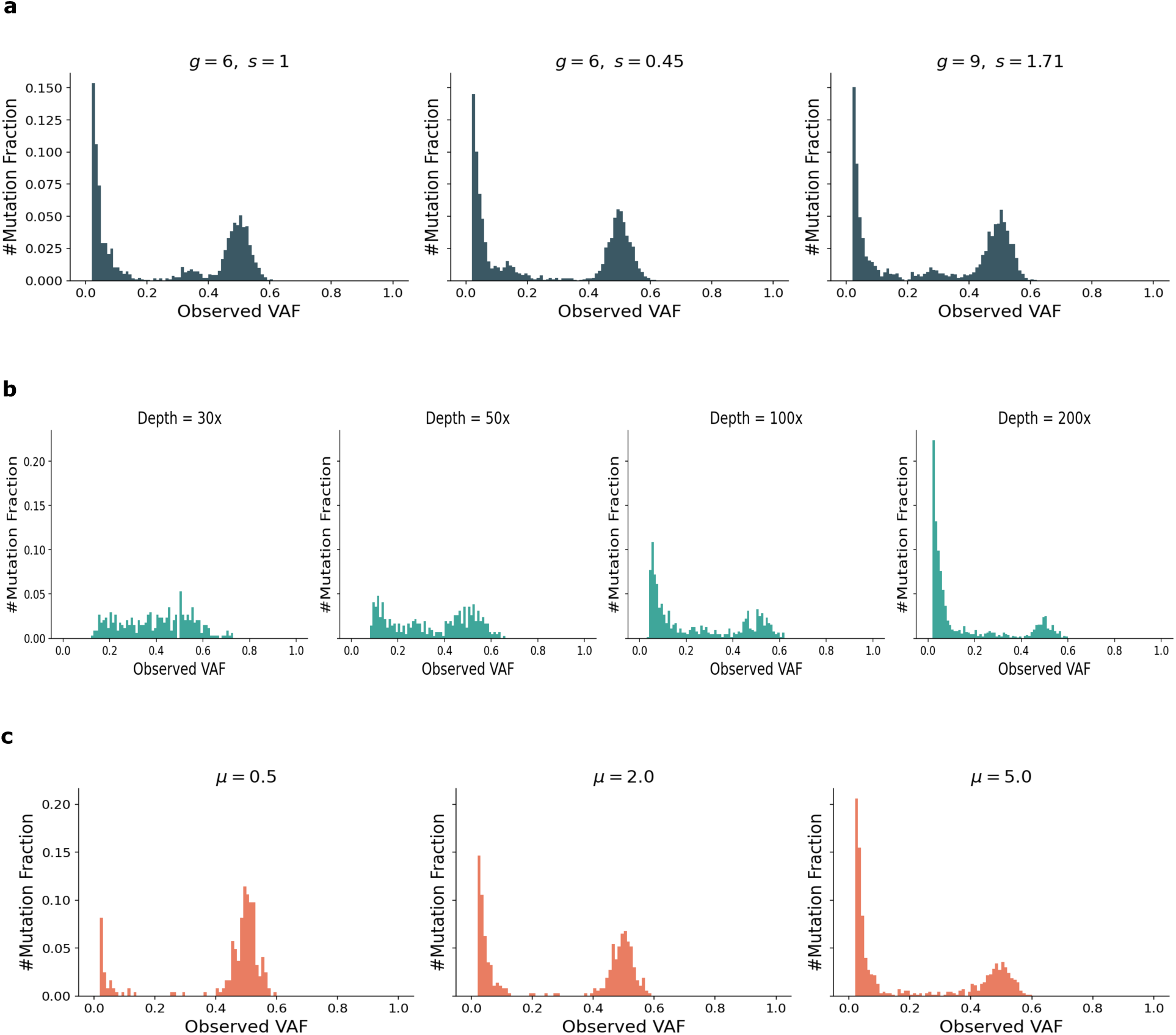
Variation in the allele frequency spectrum. **(a)** The location of the subclone cluster varies with the fraction of cells in the final population that belong to the subclone, which is in turn determined by the choice of parameters (*g, s*). **(b)** Observed allele frequency spectrum at different sequencing depths for a single simulation. **(c)** The relative sizes of the neutral tail and clonal cluster depends on the passenger mutation rates and the number of clonal mutations in the founder cell.

**Supplementary Figure S2:**
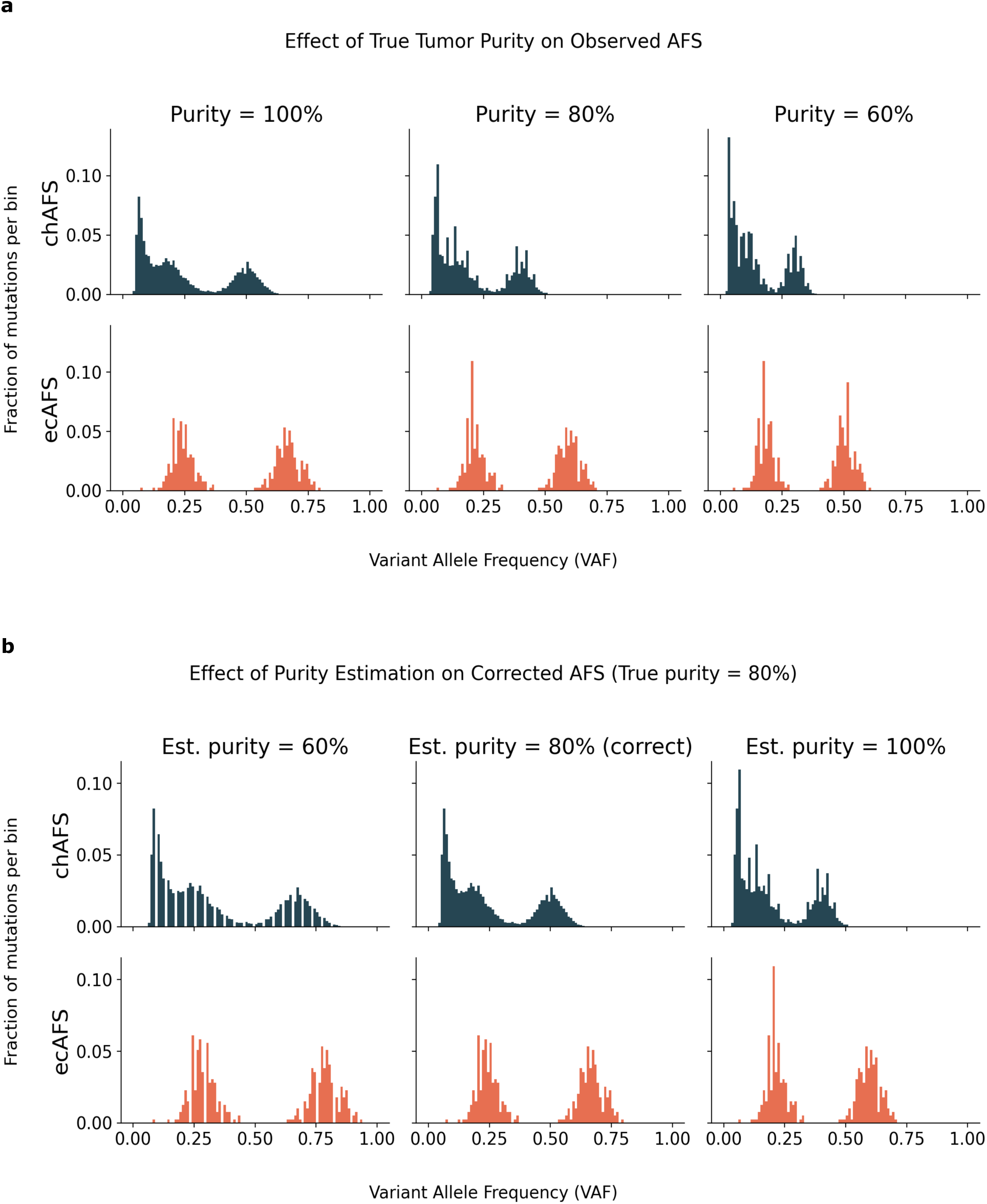
Effect of tumor purity and purity misestimation on the allele–frequency spectrum (AFS). **(a)** Observed AFS generated at three true tumor purities (100%, 80%, 60%) using a copy-number–aware dilution model. Lower purity compresses VAFs toward zero in both the chromosomal (chAFS) and ecDNA (ecAFS) regions. **(b)** AFS obtained after correcting bulk data generated at a true purity of 80% using different assumed purities (60%, 80%, 100%). Under- and over-estimation systematically distort clonal, subclonal, and ecDNA components of the AFS.

**Supplementary Figure S3:**
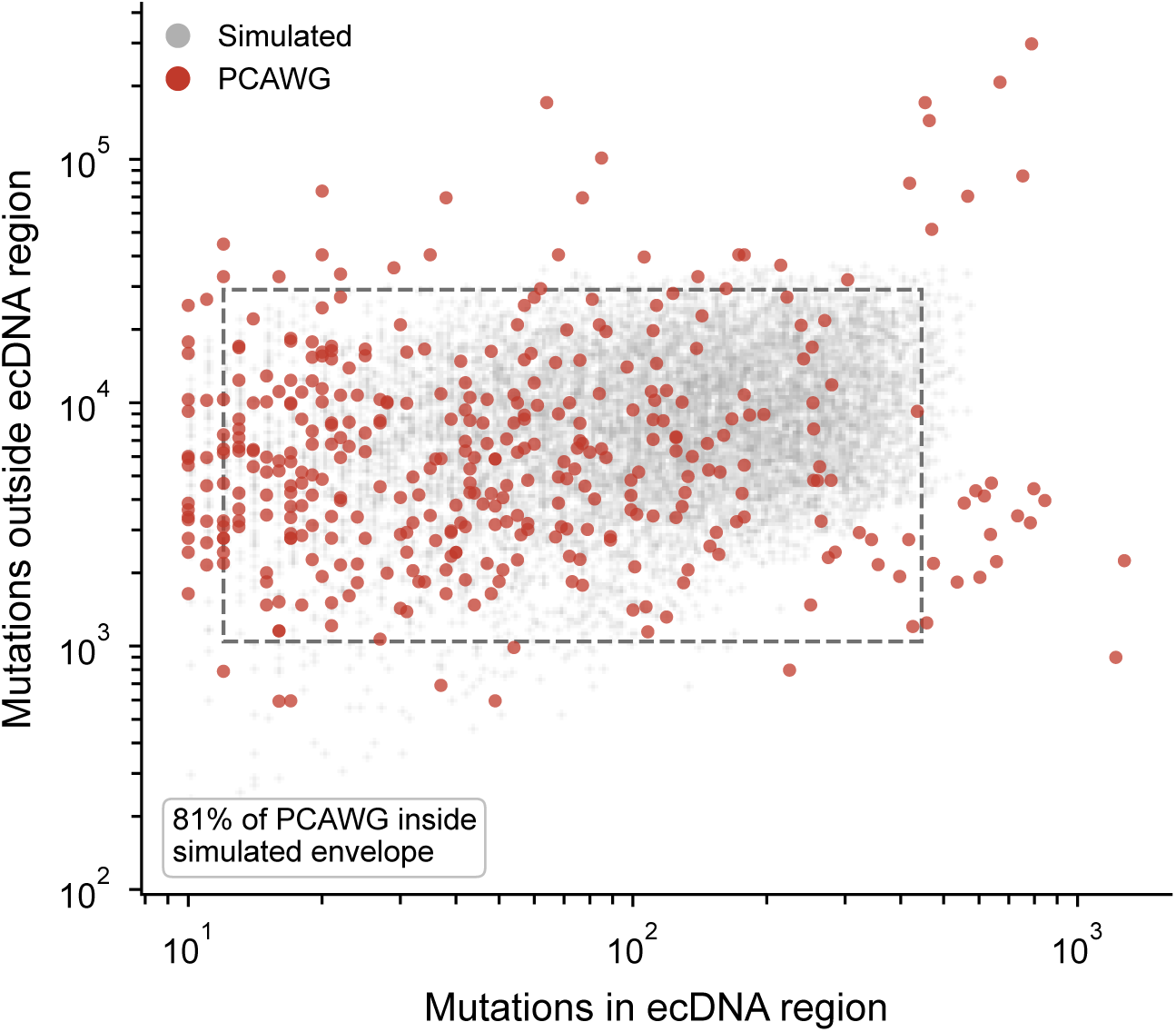
Simulated spectra span the mutation burdens observed in PCAWG tumors. Number of somatic mutations contributing to the ecDNA-region spectrum (horizontal) and to the spectrum elsewhere in the genome (vertical), for the 385 scored PCAWG ecDNA features (red) over simulated tumors pooled across sequencing depths (grey). Both sets use the same inclusion criterion of at least ten mutations in the ecDNA region. The dashed box marks the 1st–99th percentile envelope of the simulated distribution, which contains 81% of PCAWG samples; those falling outside are predominantly tumors with the highest mutation burdens. Simulation parameters are drawn from deliberately wide priors so that training data covers all parameter ranges rather than reproducing the marginal distribution of any one cohort.

**Supplementary Figure S4:**
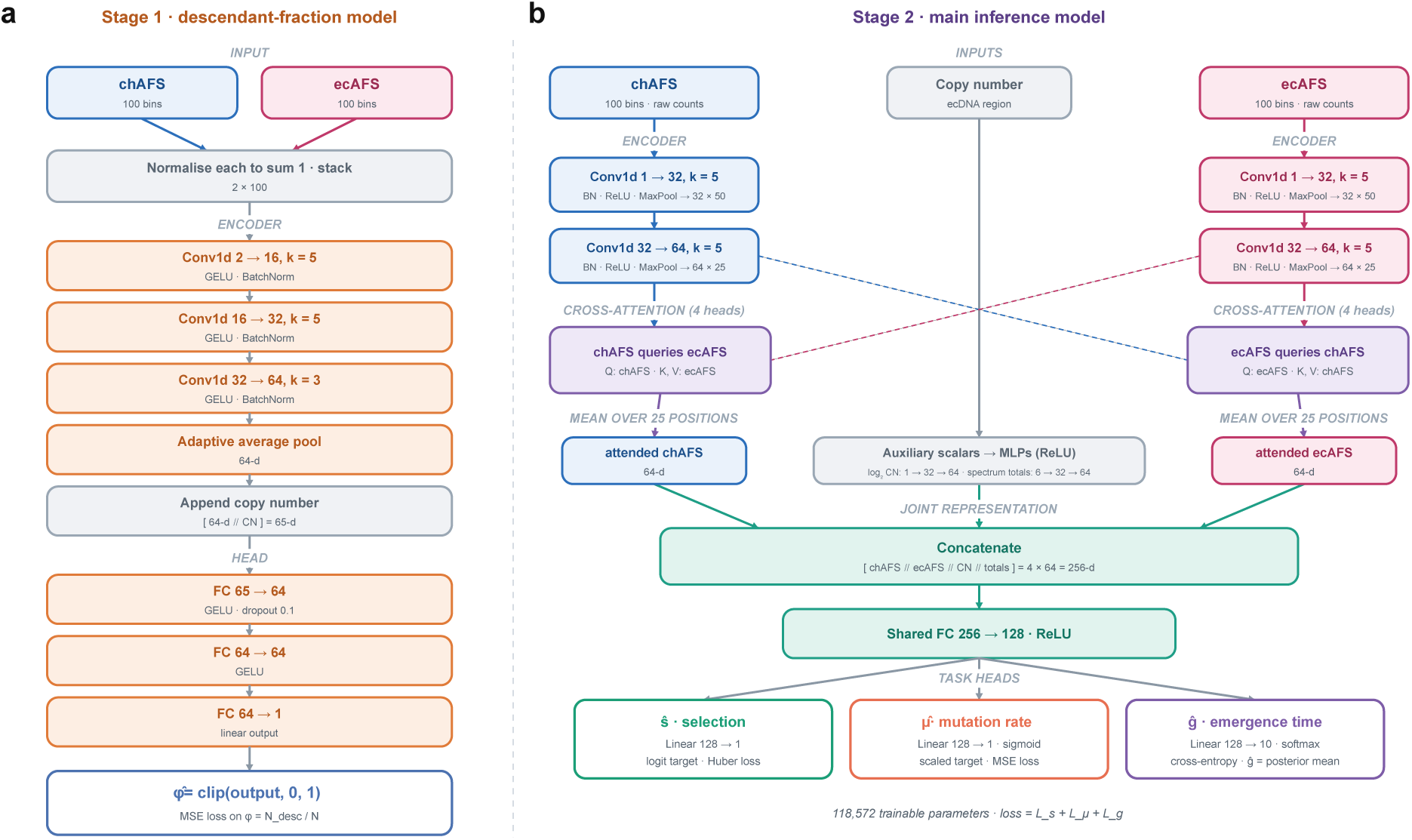
Network architectures. **(a)** Descendant-fraction model. The chromosomal and ecDNA allele-frequency spectra (chAFS, ecAFS) are each normalised to sum to one and stacked as a 2 × 100 input. Three Conv1d layers (2 → 16 → 32 → 64 channels; kernel sizes 5, 5, 3; GELU and batch normalisation) and adaptive average pooling give a 64-dimensional embedding, to which the ecDNA-region copy number is appended. Two fully connected layers (64 units, GELU, dropout 0.1) and a linear output give *ϕ̂*, trained with mean squared error against *ϕ* = *N*_desc_*/N* and clipped to [0, 1] at inference. **(b)** Main inference model. Each spectrum, as raw counts, is encoded by two convolutional blocks (kernel size 5; batch normalisation, ReLU and max pooling; 1 → 32 → 64 channels) into 25 positions of 64-dimensional features. Bidirectional cross-attention with four heads lets each spectrum query the other, and outputs are averaged over positions. The two attended 64-dimensional vectors are concatenated with 64-dimensional embeddings of the log_2_ ecDNA-region copy number and spectrum totals (256 dimensions), reduced to 128 by a shared layer, and passed to three linear heads: *s* (logistic-transformed target, Huber loss), *µ* (sigmoid, mean squared error) and *g* (softmax over ten classes, cross-entropy; *ĝ* is the posterior mean). The model has 118,572 trainable parameters.

**Supplementary Figure S5:**
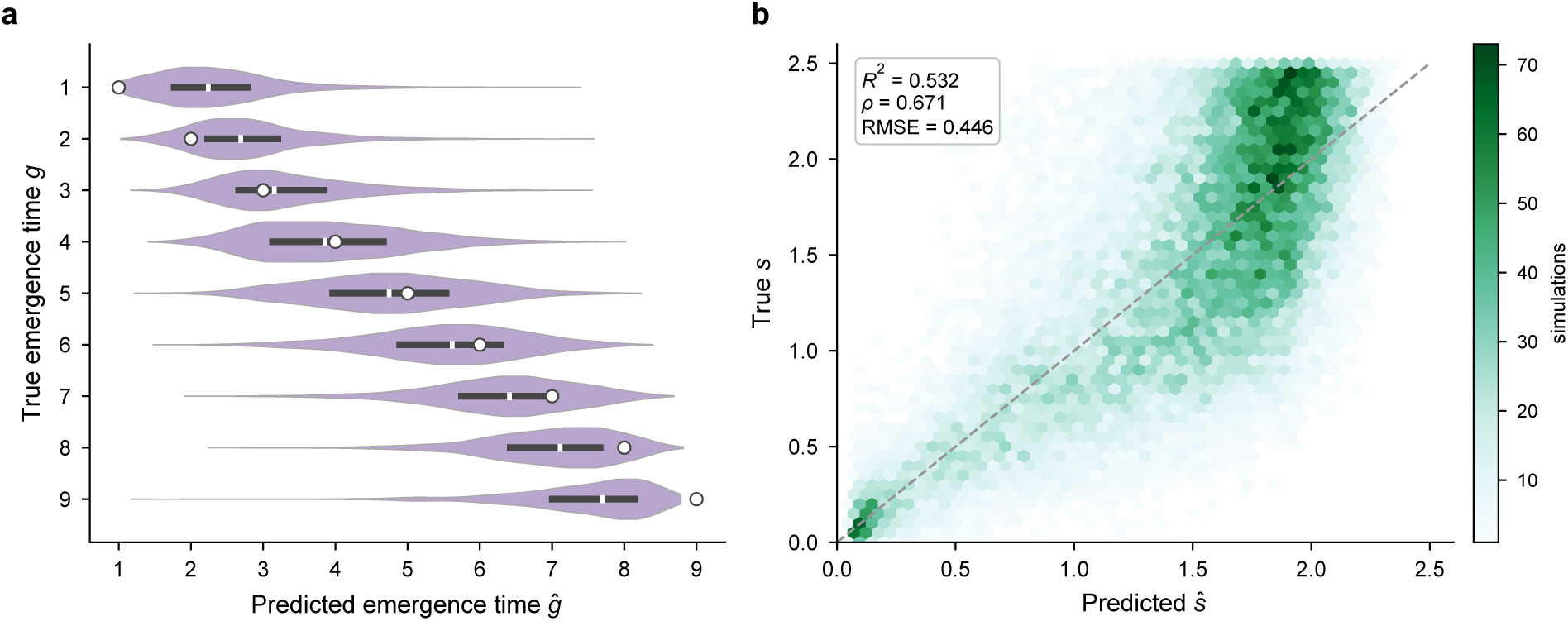
Recovery of emergence time and selection on held-out simulations (100× depth). **(a)** Predicted emergence time *ĝ* (posterior mean) for each true emergence time *g*. Violins show the density of predictions, black bars the interquartile range, and white circles the true value. **(b)** Predicted *ŝ* versus true *s* (*n* = 26,559); colour gives the number of simulations per hexagon, and the dashed line is the identity (*R*^2^ = 0.532, Spearman *ρ* = 0.671, RMSE = 0.446). Estimates saturate near *ŝ* ≈ 2 for strongly selected tumours.

**Supplementary Figure S6:**
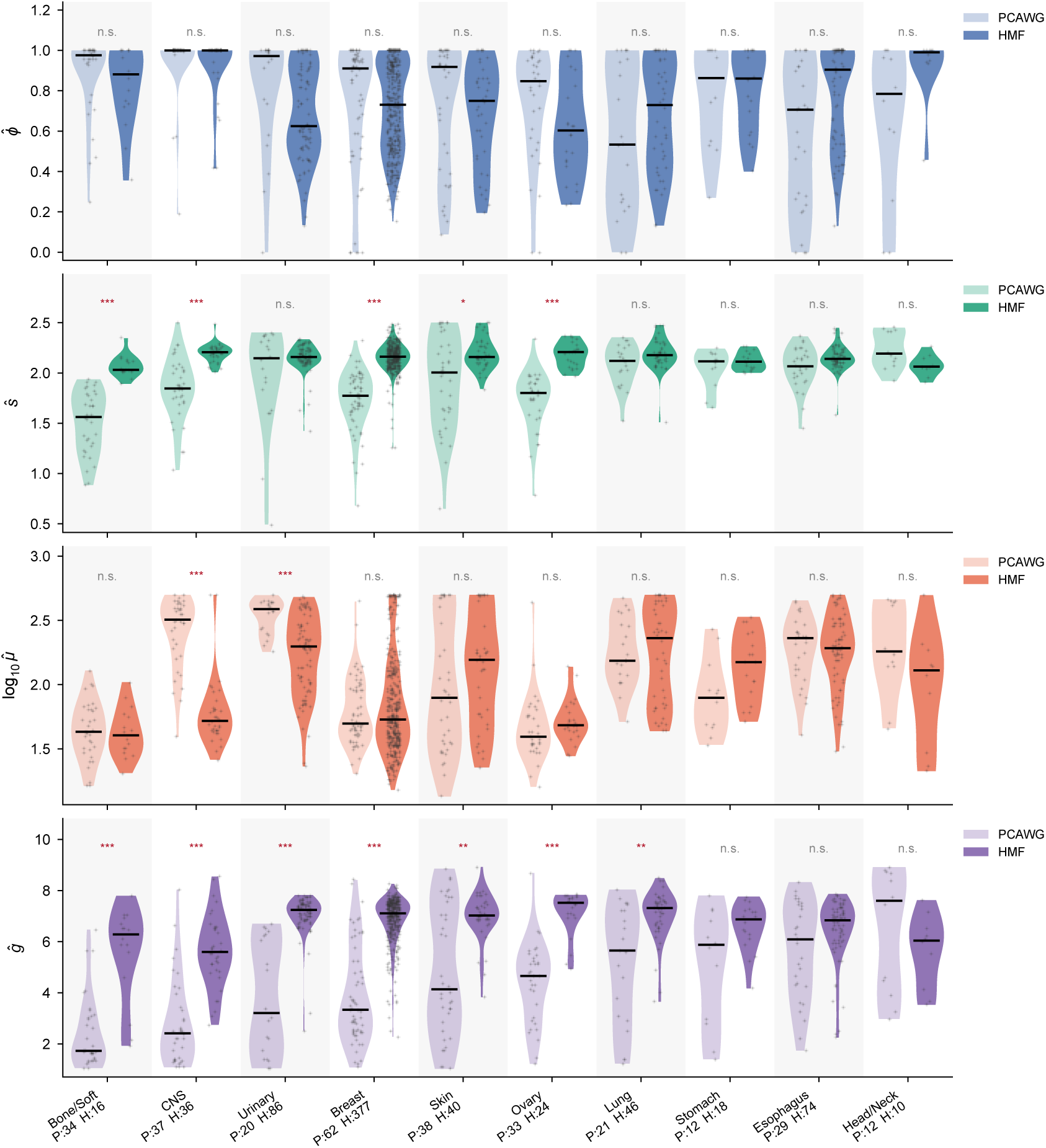
Inferred parameters by cancer type in PCAWG and HMF. Distributions of descendant fraction *ϕ̂*, selection *ŝ*, mutation rate *µ̂* (log_10_) and emergence time *ĝ* for the ten cancer groups matched between cohorts (PCAWG, light; HMF, dark), ordered by PCAWG median emergence time. Points are scored ecDNA features, horizontal bars are medians, and counts give the number of features per cohort (P, PCAWG; H, HMF). Symbols compare cohorts within each group (Mann–Whitney *U*, Benjamini–Hochberg adjusted across groups; \**P <* 0.05, \*\**P <* 0.01, \*\*\**P <* 0.001; n.s., not significant).

**Supplementary Figure S7:**
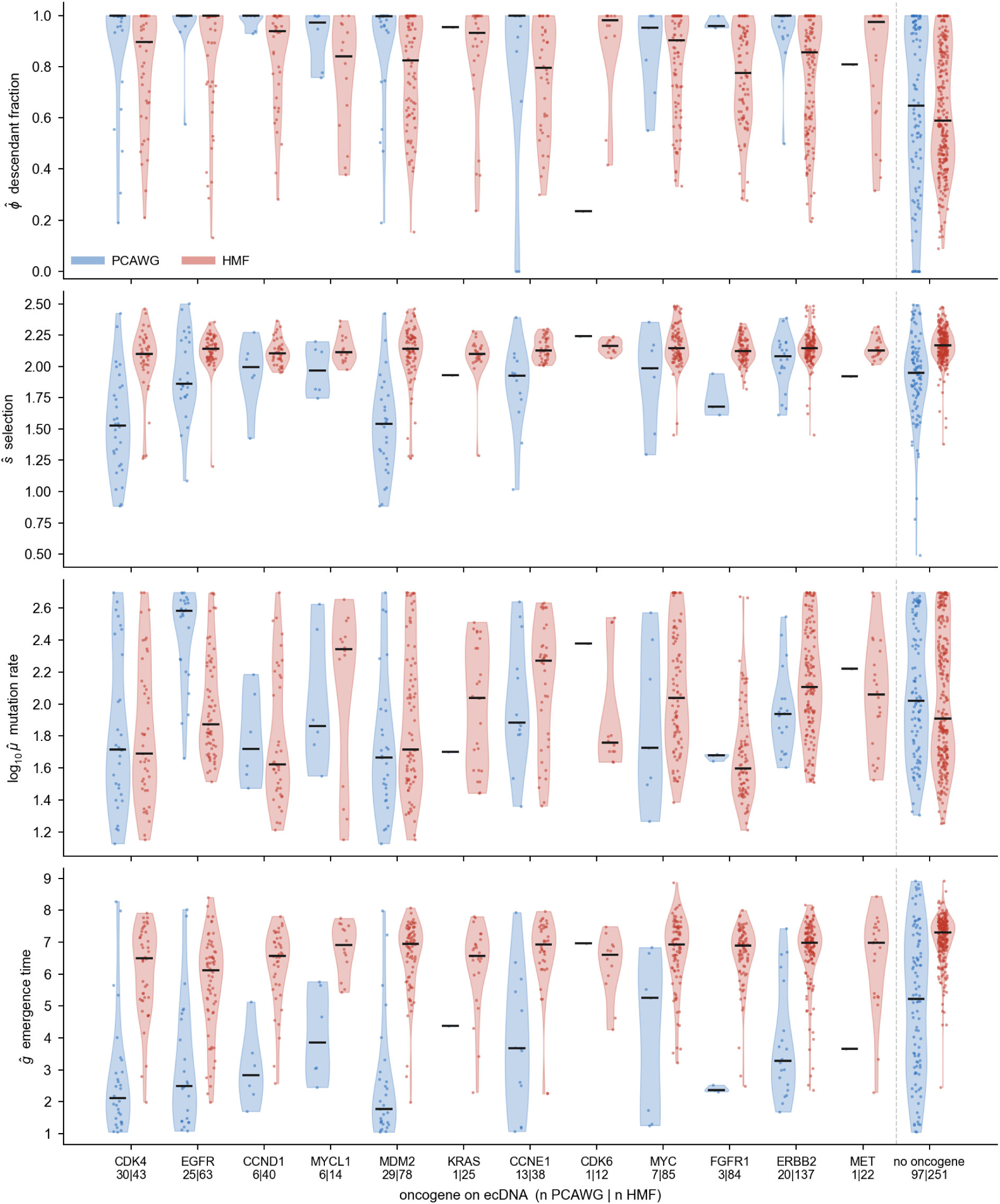
Estimated parameters for samples with matched oncogenes in PCAWG and HMF. The number of samples with ecDNA amplifying specific oncogenes is provided on x-axis for PCAWG(left) and HMF(right). While many oncogenes were amplified on ecDNA in both samples, the estimated parameters were more associated with the tumour status rather than specific oncogenes. In both cohorts, ecDNA with no known oncogenes emerged later.

**Supplementary Figure S8:**
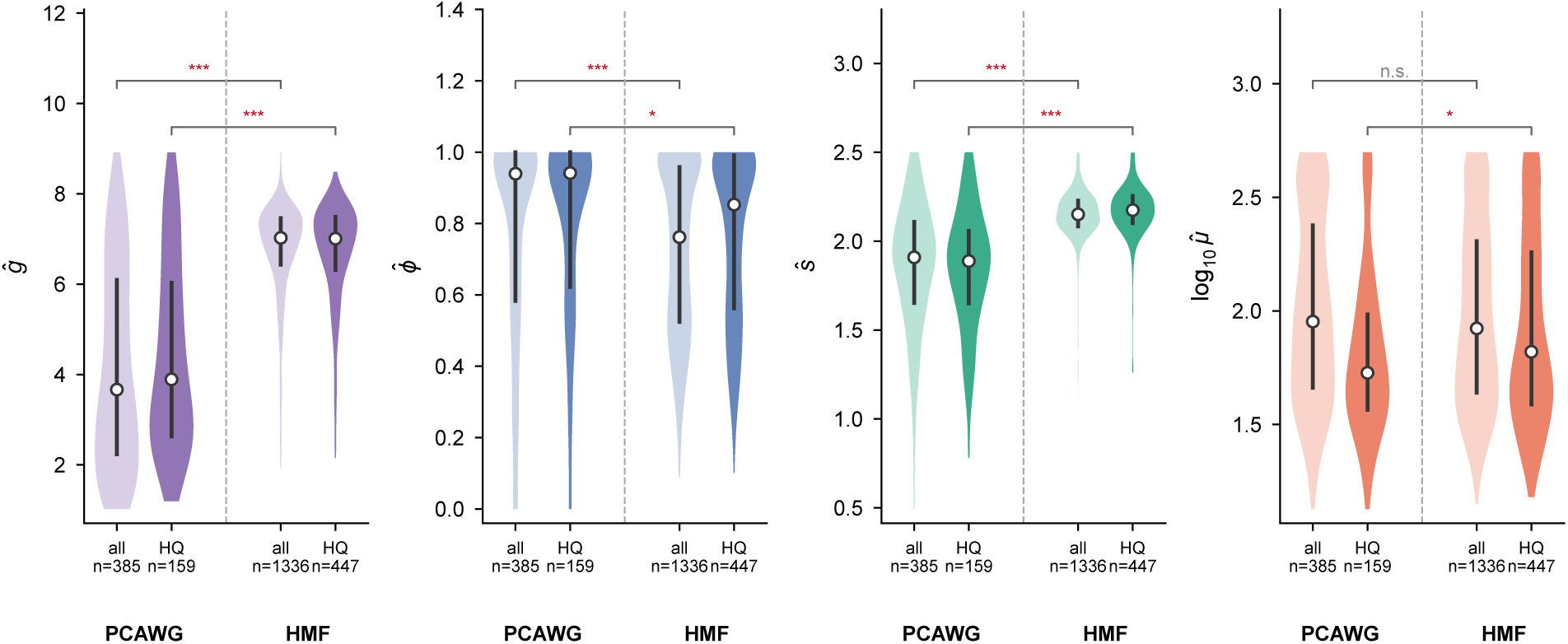
Inter-cohort differences persist in high-quality samples. Distributions of the four inferred parameters in all scored ecDNA features of PCAWG and HMF (*n* = 385 and 1,336) and in the high-quality subsets (HQ; mean sequencing depth ≥ 50× and tumour purity ≥ 0.60, the lower bound of the purity range used for training augmentation; *n* = 159 and 447). Violins show the full distribution with the median (white point) and interquartile range (bar). Brackets give two-sided Mann-Whitney *U* tests between cohorts for the full sets (outer) and the HQ subsets (inner; \**P <* 0.05, \*\**P <* 0.01, \*\*\**P <* 0.001).

## Supplementary Tables

**Supplementary Table S1:** Inference performance across sequencing depths on held-out simulated data.

| Parameter | Depth | $R^2$ | MAE |
| --- | --- | --- | --- |
| $g$ | 30× | 0.569 | 1.12 |
|  | 50× | 0.627 | 1.04 |
|  | 100× | 0.686 | 0.95 |
| $\mu$ | 30× | 0.907 | 33.1 |
|  | 50× | 0.945 | 25.3 |
|  | 100× | 0.973 | 17.3 |
| $s$ | 30× | 0.464 | 0.382 |
|  | 50× | 0.500 | 0.368 |
|  | 100× | 0.532 | 0.355 |

**Supplementary Table S2:** Positive predictive value (PPV) of predicted selection categories across sequencing depths.

| Depth | $\hat{s} > 1.25 : \Pr(s > 1)$ | $\hat{s} < 0.75 : \Pr(s < 1)$ |
| --- | --- | --- |
| 30× | 0.886 | 0.897 |
| 50× | 0.892 | 0.928 |
| 100× | 0.899 | 0.930 |

**Supplementary Table S3:**
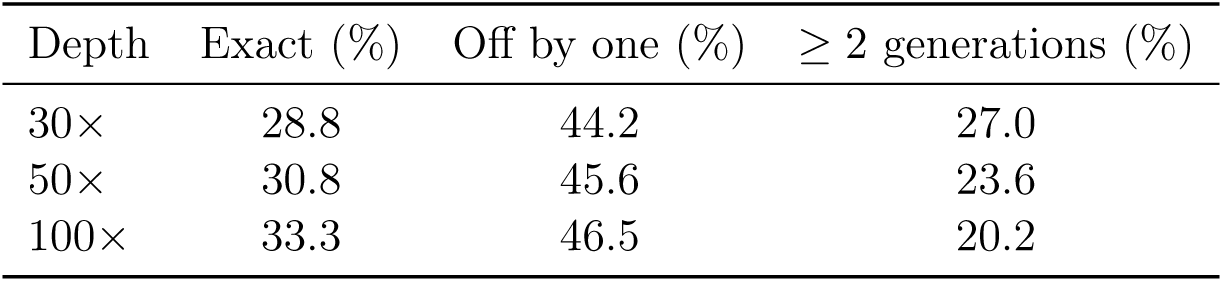
Accuracy of ecDNA emergence time inference after rounding predictions to the nearest generation. Columns are mutually exclusive and sum to 100%.

**Supplementary Table S4:** Ablation experiments on simulated data (100× depth), held-out test set. All arms were trained with seed 42. Selection is summarized by Spearman *ρ*, mutation rate by RMSE and *R*^2^, and emergence time by MAE and exact / off-by-one accuracy. The final column is Spearman *ρ*(*s*) on a purity-perturbed test set. For reference, retraining the selected architecture with ten seeds gave *ρ*_S_(*s*) = 0.669 ± 0.004 (mean ± s.d., range 0.662–0.673). Emergence-time accuracies here are computed from the most probable class; the depth-stratified tables report the rounded posterior mean.

| Model | $\rho_S(s)$ | RMSE$_{\mu}$ | $R^2_{\mu}$ | MAE$_g$ | Acc$_g^{\text{exact}}$ | Acc$_g^{\pm 1}$ | $\rho_S(s)$ pert. |
| --- | --- | --- | --- | --- | --- | --- | --- |
| Cross-attention, pooled (selected) | 0.670 | 24.5 | 0.972 | 0.96 | 0.348 | 0.785 | 0.665 |
| Cross-attention, flatten | 0.665 | 26.2 | 0.967 | 0.94 | 0.351 | 0.791 | 0.660 |
| No attention | 0.650 | 25.7 | 0.969 | 0.97 | 0.341 | 0.779 | 0.643 |
| Two-channel CNN | 0.643 | 27.2 | 0.965 | 1.00 | 0.333 | 0.765 | 0.637 |
| chAFS only | 0.641 | 25.1 | 0.970 | 1.05 | 0.320 | 0.757 | 0.636 |
| Single-task ($s$ only) | 0.666 | – | – | – | – | – | 0.659 |
| No purity augmentation | 0.670 | 25.2 | 0.970 | 0.93 | 0.349 | 0.799 | **0.431** |

**Supplementary Table S5:** Deep learning versus ABC inference performance on clean and purity-perturbed test sets (100× depth). Clean test set uses allele frequency spectra generated at 100% tumor purity. The perturbed test set applies the same two-step purity misestimation procedure used during training: allele frequencies are first mapped to a randomly drawn true purity *π*_true_ ∼ Unif(0.6, 1.0), then re-corrected using a perturbed estimate *π*_est_ ∈ [0.75 *π*_true_, 1.25 *π*_true_]. The cross-attention deep learning model was trained either with purity augmentation (matching the perturbation applied at test time) or without it (clean training data only). The ABC reference library was either the original training simulations (clean reference) or expanded sixfold by appending five purity-perturbed copies of each training simulation (augmented reference). The tolerance parameter *q* was selected independently for each reference type by cross-validation.

| Method | Test set | $\rho_S(s)$ | RMSE$_s$ | $R^2_{\mu}$ | RMSE$_{\mu}$ | Acc$_g^{\text{exact}}$ | Acc$_g^{\pm 1}$ |
| --- | --- | --- | --- | --- | --- | --- | --- |
| DL, purity-augmented training | Clean | 0.665 | 0.446 | 0.967 | 26.2 | 0.351 | 0.791 |
| DL, clean training | Clean | 0.670 | 0.445 | 0.970 | 25.2 | 0.349 | 0.799 |
| ABC, clean reference library | Clean | 0.603 | 0.487 | 0.931 | 38.0 | 0.290 | 0.697 |
| ABC, augmented reference library | Clean | 0.608 | 0.481 | 0.932 | 37.7 | 0.284 | 0.701 |
| DL, purity-augmented training | Perturbed | 0.660 | 0.451 | 0.966 | 26.7 | 0.346 | 0.782 |
| DL, clean training | Perturbed | 0.431 | 0.639 | 0.647 | 86.1 | 0.197 | 0.500 |
| ABC, clean reference library | Perturbed | 0.350 | 0.594 | 0.436 | 108.9 | 0.201 | 0.528 |
| ABC, augmented reference library | Perturbed | 0.578 | 0.501 | 0.899 | 46.0 | 0.259 | 0.654 |

**Supplementary Table S6:** Reproducibility of inferred parameters across ten random seeds. Intraclass correlation coefficients computed across ten networks differing only in random initialization, for each scored ecDNA feature in PCAWG (*n* = 385). “Single network” is the reliability of one network; “ten-network mean” is the reliability of the ensemble average by the Spearman-Brown relation. The descendant fraction is estimated by a single Stage 1 network and does not vary across the ensemble.

| Parameter | Single network | Ten-network mean |
| --- | --- | --- |
| $s$ | 0.657 | 0.950 |
| $g$ | 0.751 | 0.968 |
| $\mu$ | 0.871 | 0.985 |

**Supplementary Table S7:** Architecture and training hyperparameters of the main inference model. Values are those of the selected pooled cross-attention model, used for all depth-specific final models.

| Group | Parameter | Value |
| --- | --- | --- |
| Input | Spectrum bins per channel | 100 |
| Encoder | Convolutional blocks per spectrum | 2 |
|  | Channels per block | 32, 64 |
|  | Block composition | Conv1D, BatchNorm, ReLU, MaxPool |
| Attention | Type | Bidirectional cross-attention |
|  | Heads | 4 |
|  | Embedding dimension | 64 |
|  | Position aggregation | Global average pooling |
| Joint | Concatenated dimension | 256 |
| | Shared layer | 256 → 128, ReLU |
|  | Task heads | Linear, one per parameter |
|  | Total parameters | 118,572 |
| Training | Optimizer | AdamW |
| | Learning rate | $10^{-3}$ |
| | Weight decay | $10^{-3}$ |
|  | Batch size | 64 |
|  | Maximum epochs | 50 |
| | LR schedule | $\times 0.5$ after 3 epochs without improvement |
|  | Early stopping patience | 6 epochs |
|  | Gradient clipping | Unit norm |
| | Huber $\kappa$ | 1.0 |

**Supplementary Data 1. Per-ecDNA feature ecEvol estimates for PCAWG and HMF.** One row per scored ecDNA feature (PCAWG, *n* = 385; HMF, *n* = 1,336), with sample identifiers, cancer type, copy number, purity, the sequencing depth of the model used, and the inferred descendant fraction, selection coefficient, mutation rate and emergence time. Estimates are means over ten networks; standard deviations across networks and the posterior probability that *g* ≤ 2 are also given. Column definitions are on the README sheet.

